# Phosphorothioate Substitutions in RNA Structure Studied by Molecular Dynamics Simulations, QM/MM Calculations and NMR Experiments

**DOI:** 10.1101/2020.10.28.359059

**Authors:** Zhengyue Zhang, Jennifer Vögele, Klaudia Mráziková, Holger Kruse, Xiaohui Cang, Jens Wöhnert, Miroslav Krepl, Jiří Šponer

## Abstract

Phosphorothioates (PTs) are important chemical modifications of the RNA backbone where a single non-bridging oxygen of the phosphate is replaced with a sulphur atom. PT can stabilize RNAs by protecting them from hydrolysis and is commonly used as tool to explore their function. It is, however, unclear what basic physical effects PT has on RNA stability and electronic structure. Here, we present Molecular Dynamics (MD) simulations, quantum mechanical (QM) calculations, and NMR spectroscopy measurements, exploring the effects of PT modifications in the structural context of the Neomycin-sensing riboswitch (NSR). The NSR is the smallest biologically functional riboswitch with a well-defined structure stabilized by a U-turn motif. Three of the signature interactions of the U-turn; an H-bond, an anion-π interaction and a potassium binding site; are formed by RNA phosphates, making the NSR an ideal model for studying how PT affects RNA structure and dynamics. By comparing with high-level QM calculations, we reveal the distinct physical properties of the individual interactions facilitated by the PT. The sulphur substitution, besides weakening the direct H-bond interaction, reduces the directionality of H-bonding while increasing its dispersion and induction components. It also reduces the induction and increases dispersion component of the anion-π stacking. The sulphur force-field parameters commonly employed in the literature do not reflect these distinctions, leading to unsatisfactory description of PT in simulations of the NSR. We show that it is not possible to accurately describe the PT interactions using one universal set of van der Waals sulphur parameters and provide suggestions for improving the force-field performance.

## Introduction

The chemical modifications of RNA can have important and diverse effects on RNA function, gene transcription and translation regulation.^1^ The modified chemical groups can, for example, change the RNAs expression rate, alter its stability by inhibiting the hydrolysis by nucleases or by modulating the interactions between RNA and other molecules.^2^ In RNA molecules, the thio modification refers to a chemical modification in which sulphur substitutes oxygen.^3–12^ If the substitution occurs on a nonbridging oxygen of a phosphate group, a phosphorothioate (PT) is formed. The PT modification in RNA is often artificially introduced as a tool of biochemical research.^11^ However, very recently it was also reported to occur naturally in both prokaryotic and eukaryotic RNAs.^10^ The PT can alter the biochemical properties of nucleic acids and their interactions.^5^ In siRNA, PT increases or decreases gene silencing by adjusting the siRNA stability.^6^ It can also influence the binding of proteins and metal ions to nucleic acids.^7–9^ PT modifications of mRNA can also increase its translational activity.^13^ A significant hurdle for experimental studies of PT is the presence of two non-bridging oxygens in every RNA phosphate group as potential targets for a thio modification, leading to two PT enantiomers; Rp and Sp; as shown in Figure 1A.^8,10,12^ The two enantiomers can for instance influence ribozyme reaction profiles in very different ways.^14^ Separate synthesis of the two enantiomers or purification of their mixtures are non-trivial issues and subject of ongoing experimental research.^15–17^ On the other hand, the PT enantiomer can be unambiguously selected in theoretical studies. Despite their significant biological effects on RNA metabolism, the structural and dynamical impacts of PT modifications are poorly understood.^18^

**Figure 1:**
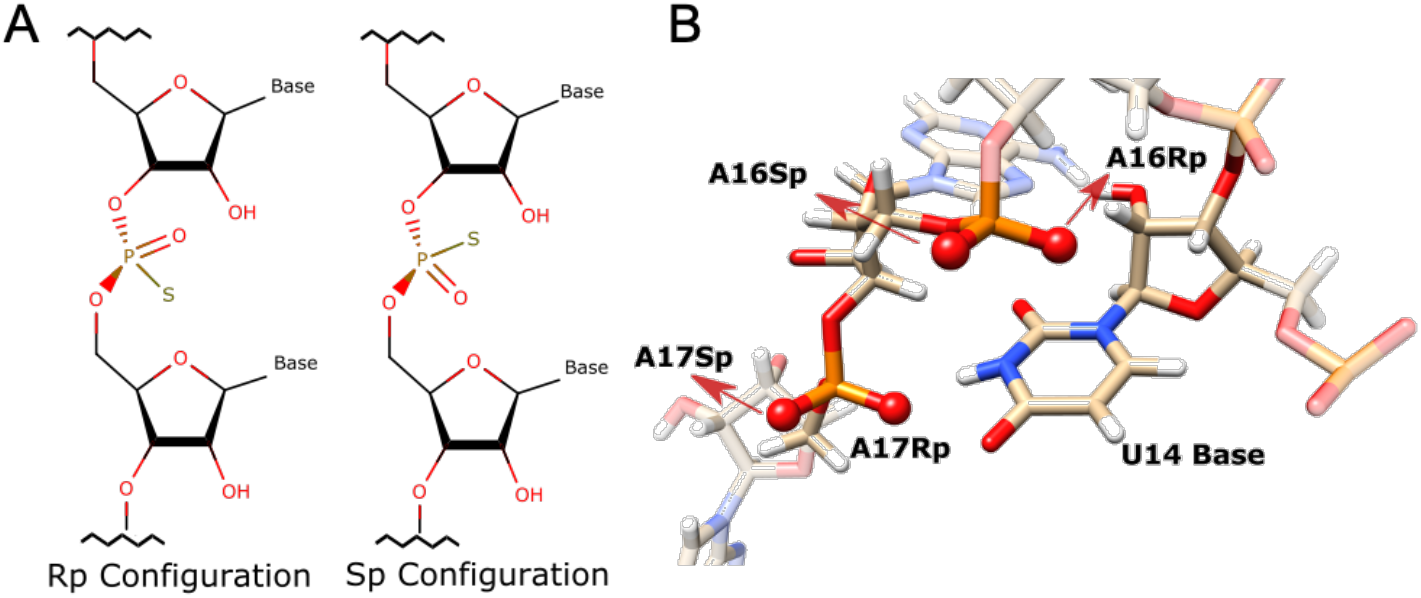
A) Scheme of the two PT enantiomers where sulphur replaces single non-bridging oxygen in RNA phosphate linkage. B) The stereochemistry of the A16 and A17 phosphate linkages in the context of the NSR – the model system used in this work.

Here, we present a theoretical study of the influence of PT modifications on RNA structure and dynamics using MD simulations and QM/MM and QM calculations, supported by NMR spectroscopy experiments. MD is a computational method which applies a carefully calibrated set of molecular mechanical (MM) empirical potentials (force fields) to describe motions of molecules at the atomistic level.^19,20^ It can explore molecular motions with high spatial and temporal resolution, unrivalled by any other method. However, the MD simulations must be carefully interpreted, as the force fields are collections of parametrized terms which, despite looking at first sight very intuitive, are entirely unphysical.^20,21^ In contrast, quantum mechanics (QM) calculations may provide, in principle, physically correct description of the studied systems. However, the large computational demands of QM limit the size of the studied systems and prohibit Boltzmann sampling of microstates, which is absolutely crucial for a meaningful description of biochemical systems. The QM and MM approaches can be combined into hybrid (QM/MM) calculations, in which the core molecular region of interest is described by QM and the remaining parts of the system by MM.^22,23^ QM/MM allows to circumvent the system size limitations but Boltzmann sampling is still inaccessible. Both QM and QM/MM calculations can help validate force fields used by MD simulations.^24–26^

The RNA molecule utilized for our study of PT is the Neomycin-sensing riboswitch (NSR), a widely studied riboswitch containing only 27 nucleotides.^27–30^ In general, riboswitches are RNA regulatory elements, commonly located in the 5′-untranslated regions of mRNAs. They regulate translation by changing their conformation upon binding of a ligand, such as a metabolite, metal ion, or antibiotics.^20,31–34^ The NSR is the smallest known biologically functional riboswitch. Upon binding aminoglycosides like neomycin or ribostamycin, it forms a stable hairpin within the 5′-UTR region of the mRNA, blocking translation initiation downstream of the riboswitch. In the ligand-bound state the NSR has a hairpin-like structure with an A-form helix from base pair G1:C27 to U13:U18 and an apical loop from U14 to A17 (Figure 2).^28^ The A-form helix is interrupted with a bulge on one strand, from C6 to U8. The apical loop U14-A17 segment and the closing U13:U18 base pair together form the 5′-UNR-3′ (N – any nucleotide, R – purine) U-turn, which is a recurrent RNA motif commonly capping A-RNA helices.^35,36^ The bulged-out A17 nucleotide is an NSR-specific insertion into the U-turn loop which facilitates contact with the antibiotics, bound below the apical loop. Structural details of the NSR and its U-turn motif were unveiled by previous studies.^28,29^ The U-turn of the NSR is stabilized by two signature H-bonds between U14(N3)-A17(OP2) and U14(O2’)-A16(N7), respectively. We will refer to these two hydrogen bonds as HB1 and HB2 in the course of this work. There is also an anion-π interaction between the U14 base and the A16(OP2) atom. The anion-π interaction is a weak non-bonded interaction^37^ that was nevertheless suggested to contribute to the stability of the U-turn motif.^36^ Lastly, there is a highly-occupied monovalent cation binding site near the U14(O4) atom which was recently identified by MD simulations and NMR titrations as a crucial and evolutionary conserved element of the U-turn motif.^29^ Although the first uracil of the 5′-UNR-3′ U-turn motif is regarded as essential and conserved, U-turns can also have a cytosine at this position.^29,38^ In this case, the cytosine is N3-protonated to maintain the signature H-bond HB1 while the overall positive charge of the protonated base substitutes for the cation binding site.^29^

**Figure 2:**
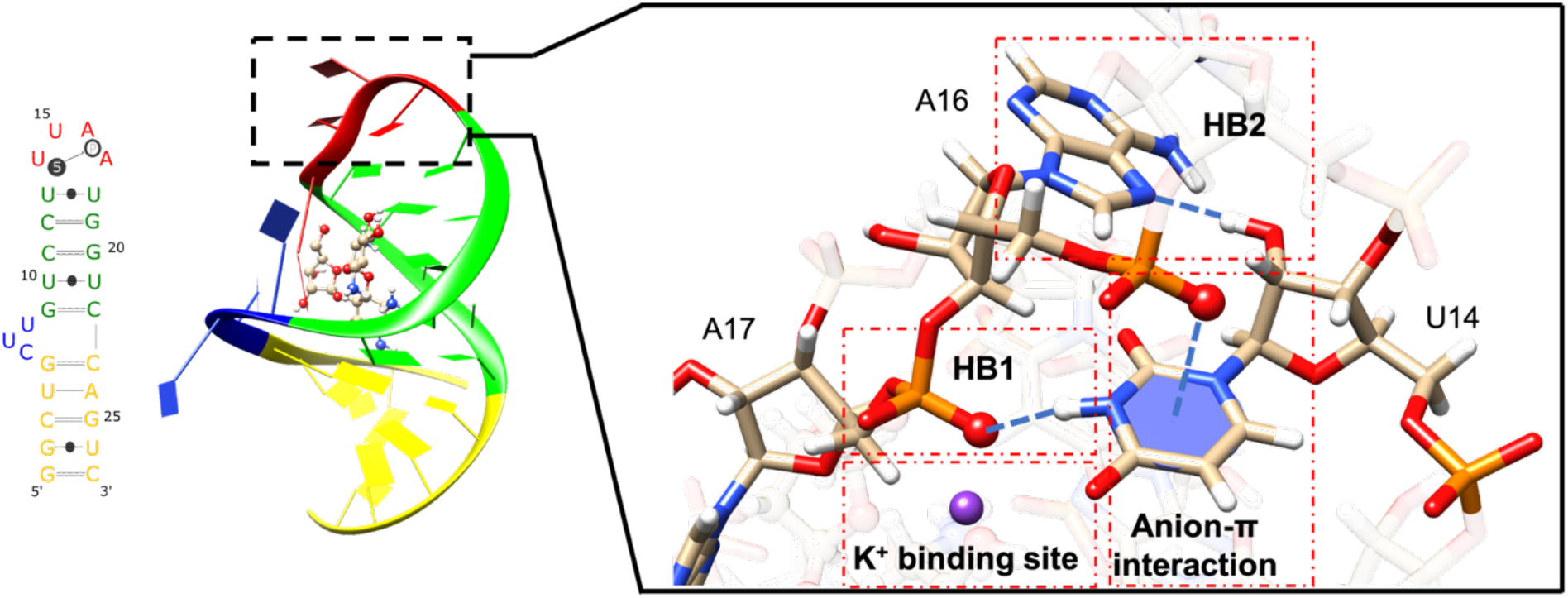
Secondary structure of the NSR and its 3D structure with the bound ribostamycin shown in CPK representation. The A-RNA helical segments are shown in green and yellow, respectively, separated by the flexible bulge in blue. The U-turn loop is in red. Its two signature H-bonds (termed as HB1 and HB2, respectively), the anion-π interaction, and the potassium binding site are shown in detail. The Rp oxygen atoms of the A16 and A17 phosphate groups are highlighted as spheres.

We have investigated the effect of PT modification on the NSR by introducing single PT modifications in either A16 or A17 phosphate groups. The U-turn signature interactions mentioned above involve the phosphate groups of A16 and A17 and could therefore be affected by these PTs (Figure 1B and Figure 2). Namely, PT modifications with the Rp configuration at A17 and A16 should directly influence the HB1 signature H-bond and the anion-π interaction, respectively (Figure 2). PTs can also alter solute-solvent interactions, namely the potassium binding site near U14(O4)^29^ and interactions with water. We carried out NMR spectroscopy, MD simulations, QM/MM optimizations and interaction energy QM scans and describe the changes in structure, dynamics and stability of NSR resulting from PT modifications. Our results provide new insights into the effects of PT modifications in RNA in context of different molecular interactions which could provide a reference for interpreting future experimental studies involving PT modifications. Previously, it was shown that the wild-type NSR is well described by contemporary force fields and can be a useful system for force-field testing and computational modeling studies.^29^ Therefore, we also evaluate the quality of MM description of the PT modification by comparing MM data with reference QM and QM/MM computations. The QM data explain the physical nature of the unique interaction capabilities of the PT compared to the phosphate. We also demonstrate that it is not possible to accurately describe PT interactions in MD using one universal set of sulphur van der Waals parameters.

## Methods and Material

### MD Simulations

The structure of the NSR complexed with ribostamycin determined by high-resolution solution NMR (PDB ID: 2n0j)^28,30^ was utilized as starting structure in all MD simulations. To study the effects of the PTs in A16 and A17 phosphates, and to examine their effect when U14 is replaced with C14^+^, we prepared simulation systems as shown in Table 1. The starting systems with PT modifications and the C14^+^ substitution were prepared by molecular modeling using the available wild-type experimental NSR structure 2n0j as a template. We used the bsc0χOL339,40 (OL3) RNA force field as implemented in AMBER18.^41^ The GAFF force field was used to describe the ribostamycin ligand.^42^ To obtain partial charges of the PT group, we calculated the electrostatic potential of dimethyl-thiophosphate (DMTP) in Gaussian 09 (rev. A02)^43^ using the HF/6-31G* level of theory. This was followed by partial-charge fitting and residue assignment of the whole thiophosphate group in antechamber via the RESP^44,45^ procedure. Parameters for the ribostamycin ligand and for protonated cytosine C14^+^ were taken from earlier simulation study of the NSR.^29^ Each system was processed by a series of minimizations and equilibrations as described in the Supporting information. The reference wild-type and mutC14 simulations with no chemical modifications had simulation times of 10 microseconds, while the thio-modified systems were simulated in four parallel simulations, each one-microsecond long (SI Appendix Table S1). All MD simulations were performed in truncated octahedron boxes of SPC/E^46^ water molecules along with 0.15 M concentration of KCl^47^ ions. In our earlier simulation study of NSR, we reported excellent performance of this specific combination of OL3 RNA force field with the water model and ions and demonstrated that NSR simulations are not sensitive to the choice of water model.^29^ The shortest distance from the solute to the periodic box edge was 12 Å. In selected simulations, we have used the structure-specific HBfix potential^48^ to increase stability of the HB2 signature H-bond (see Supporting Information for further details).

**Table 1:** List and designations of the studied systems.^a^

| PT configuration | A16 PT |  | A17 PT |  |
| --- | --- | --- | --- | --- |
|  | with U14 | with C14<sup>+</sup> | with U14 | with C14<sup>+</sup> |
| **Rp** | A16Rp/U14 | A16Rp/C14 | A17Rp/U14 | A17Rp/C14 |
| **Sp** | A16Sp/U14 | A16Sp/C14 | A17Sp/U14 | A17Sp/C14 |
| **No PT** | with U14 |  | with C14<sup>+</sup> |  |
|  | wt |  | mutC14 |  |

<sup>a</sup> Depending on stereochemistry of PT, and on whether U14 or C14<sup>+</sup> is present, there are ten different NSR systems in total – eight PT systems and two without PT. The items in the table are designations of the individual NSR systems as utilized throughout this study.

### QM/MM Optimizations

As starting structures for non-periodic QM/MM calculations in a water droplet, we manually selected snapshots from the MD simulations with a geometry that is most representative of the common simulation trends. For C14^+^ systems, where we studied bifurcation of the HB1 interaction, we selected simulation snapshots where such bifurcation occurred. As the MD simulations were done in periodic boundary conditions, we only took the solute coordinates from the simulation snapshots. In addition, coordinates of the potassium ion near the U14(O4) atom were used for the U14 systems. We then added a sphere of water molecules with radius of 34 Å along with enough potassium ions to neutralize the system. The added potassium ions were kept at least 5 Å away from the solute so they would not bias the QM electron densities. This approach was developed and validated in Ref^26^. Afterwards, we performed 2000 cycles of MM minimization and an equilibration run for 0.5 ns, while the solute and the single potassium near U14(O4) were frozen. Atoms ranging from C11(C4′) to U21(C5′) were selected as the QM region, encompassing the entire U-turn motif. In U14 systems, the potassium near U14(O4) atom was also included. The QM region corresponded to 317 and 318 atoms in U14 and C14^+^ systems, respectively. After that, we used the sander program of AMBER16^49^ as the interface of Turbomole^50,51^ to perform non-periodic QM/MM optimizations. The composite DFT method PBEh-3c^52^ was used to describe the QM region (Table 2) while MM parts of the systems were described by the same force fields as in MD. The electrostatic interaction between MM point charges and QM atoms was solved by electrostatic embedding scheme in which the MM region can polarize the QM electron density. Explicit link atoms were used to handle charges at the covalent boundary between the QM and MM regions.^23^ To compare QM/MM and MM results, equivalent MM optimizations were also carried out for each system.

**Table 2:** List of utilized QM methods.

| QM Method | Basis set | Application |
| --- | --- | --- |
| PBEh-3c | def2-mSVP | QM/MM optimization, interaction energy scan |
| SAPT2+(3) δMP2 | jun-cc-pVTZ | Interaction energy decomposition |
| DLPNO-CCSD(T) | aug-cc-pV(T-Q)Z | Interaction energy scans validation |

### Interaction Energy QM and MM Scans

Interaction energy scans performed on small model systems representing the signature interactions of the U-turn were used to further analyze the effects of the PTs. The initial coordinates for the models were excised from the QM/MM optimized structures. The atoms selected for the models included the U14 base along with either the A17 or A16 thiophosphate/phosphate for HB1 or anion-π interaction models, respectively. The C5′ and C3′ carbons connected to these thiophosphates/phosphates and the C1′ carbon connected to the U14 base were also included in the models and hydrogens were added to them, forming dimethyl-thiophosphate/phosphate (DMTP/DMP) and methyl-uracil (MU). The hydrogens of the methyl groups were added by Avogadro,^53^ and their coordinates were optimized by the PBEh-3c QM calculation while the other atoms were fixed. Afterwards, using an in-house program, we created a series of systems with different distances between DMTP/DMP and MU. The scanned distance ranges are detailed in the Supporting Information. For each QM distance scan calculation, we also did a corresponding distance scan calculation using MM. The PBEh-3c calculations were done by Turbomole version 7.3.^50^ In some cases, the symmetry-adapted perturbation theory (SAPT2+(3) δMP2 with jun-cc-pVTZ basis set; SAPT in the following text)^54–56^ (Table 2) was used to decompose the interaction energies into electrostatic, dispersion, exchange and induction energy components using the PSI4 program.^57^ We also used the more rigorous DLPNO-CCSD(T) method^58–60^ using ORCA V4.1.0^61,62^ to validate selected QM distance scans (see Supporting Information for details). We have conducted the distance scans both in vacuum as well as in COSMO implicit solvent model^63^ using a relative permittivity of 78.4 and the new FINE cavity construction.^64^ For MM calculations done in implicit solvent, we used Generalized-Born/Surface Area (GBSA) model.^65^ A spline function was used to interpolate the distance at which the interaction energy was at a minimum.

In addition to the DMTP/DMP-MU models, we also performed QM distance scans of DMTP/DMP-water model systems. Technical details of these calculations are described in the Supporting Information.

### Optimization of van der Waals (vdW) parameters

Both the small model systems as well as the QM/MM calculations revealed relatively poor agreement between MM and QM calculations when using the standard AMBER vdW parameters for sulphur (see below). We have subsequently re-run these MM calculations with an alternative vdW sulphur parameter from CHARMM (termed “thiolate” parameter, see Table 3). We also repeated the MD simulations using the thiolate parameter to verify its performance. In addition to globally changing the sulphur vdW parameters, we also tested an NBfix (Non-Bonded fix) implementation of the thiolate parameter in which the nonbonded interactions of the sulphur atom are tuned only between specific atomic pairs (Table 3). The NBfix approach allows solving a situation in which a single global vdW parameter is not ideal to describe non-bonded interactions with all parts of the studied system.^66^ It allows tuning the vdW parameter with potentially fewer undesired side-effects and is particularly suitable when targeting the size of the MM atom and its short-range repulsion.^67^ The simulated systems and their timescales are summarized in Table S1.

**Table 3:** Overview of the utilized sulphur (SP2) vdW parameters and methods of their implementations in calculations.

| Force field modification | Details<sup>a</sup> |
| --- | --- |
| **Original AMBER force field<sup>b</sup>** | $R(\text{SP2}) = 2.0 \text{ \AA}$, $\epsilon(\text{SP2}) = 0.25 \text{ kcal/mol}^{41}$ |
| **Thiolate parameters<sup>b</sup>** | $R(\text{SP2}) = 2.2 \text{ \AA}$, $\epsilon(\text{SP2}) = 0.47 \text{ kcal/mol}^{68}$ |
| **NBfix<sup>c</sup>** | $R(\text{SP2}) = 2.2 \text{ \AA}$, $\epsilon(\text{SP2}) = 0.47 \text{ kcal/mol}$ for SP2 - H3, N3, $O_{\text{water}}$, $K^+$, $\text{Cl}^-$ nonbonded pairs<br>$R(\text{SP2}) = 2.0 \text{ \AA}$, $\epsilon(\text{SP2}) = 0.25 \text{ kcal/mol}$ for all other nonbonded pairs |

<sup>a</sup>The $R$ and $\epsilon$ refer to the radius and well-depth, respectively, of the vdW parameters. The radii of vdW parameters in this work always refer to the $R_{\text{min}}/2$ value. <sup>b</sup>vdW parameters of sulphur were globally applied for all its nonbonded pairwise interactions. <sup>c</sup>Thiolate parameters were applied for the specified pairwise interactions while standard AMBER sulphur vdW parameters were applied for all other nonbonded pairwise interactions.

### Analyses

The visualization and analyses of simulation trajectories were done in Chimera,^69^ VMD^70^ and cpptraj.^41^ RStudio^71^ was used for statistical analyses and visualization of the extracted datasets. For hydrogen bond analysis, we considered the hydrogen bond present if the heavy atom donor-acceptor distance was lower than 4 Å and the H-bond angle was larger than 120°. The 4 Å distance cutoff was enough to accommodate the sulphur-based hydrogen bonds. The anion-π interaction was considered present when the distance between the base plane and the oxygen or the sulphur was below 3.8 Å or 4.3 Å, respectively, and the distance between the oxygen/sulphur and the perpendicularly projected point from the geometric center of the base was below 1.5 Å. Note that except for the HB1 and anion-π populations analyses, we excluded all simulation frames in which the U-turn motif underwent a temporary disruption of its key HB1 interaction for the analysis purposes. Although this could stem from a genuine dynamics of the U-turn motif on a microsecond timescale (see also Ref^29^ for more details) we excluded such parts of simulations from the analyses to prevent contamination of the datasets with statistical uncertainty. All trajectories were visually monitored to guarantee that no significant developments were missed.

### RNA sample preparation

The wt and mutC14 variants of the NSR were synthesized by *in vitro* transcription with T7 RNA polymerase. Linearized plasmid DNAs containing a 3′ - hammerhead ribozyme sequence were used as templates to obtain RNAs with uniform 3′-ends. The RNA synthesis was done using 4 mM unlabeled commercially available nucleotide triphosphates (Merck). The obtained RNAs contain a 5′-triphosphate and a 2′,3′-cyclic phosphate at their 5′- and 3′-termini, respectively, due to the cleavage by the ribozyme. *In vitro* transcription with T7 RNA polymerase in the presence of α-thio-ATP was used to obtain PT-modified NSR with an Rp configuration of the thiophosphates. In this case, we used synthetic single-stranded DNA templates with a double-stranded promotor region without a 3′ ribozyme sequence since α-thio-ATP incorporation might interfere with ribozyme activity. As a result, these RNAs can contain additional non-templated nucleotides at their 3′-ends as well as 3′-terminus with a 3′-OH group. Thus, for these RNAs, slightly different chemical shifts and/or multiple sets of NMR resonances for the first two base pairs of the NSR stem are observable in comparison to the unmodified RNAs transcribed as hammerhead ribozyme fusions. The synthesis of these RNAs was done using 4 mM cytosine-, guanosine- and uridine-5′-triphosphate and 4 mM adenosine-5′-(α-thio)-triphosphate (Jena Bioscience). Selectively ^13^C,^15^N-C labeled and PT-modified U14C RNAs were synthesized using 4 mM ^13^C,^15^N labeled cytosine-5′-triphosphate (Silantes), 4 mM adenosine-5′-(α-thio)-triphosphate and 6 mM unlabeled guanosine- and uridine-5′-triphosphate. The RNA transcripts were purified by preparative PAGE according to standard protocols, desalted via PD-10 columns (GE Healthcare) and folded into monomeric hairpin forms by heating to 95 °C for 10 minutes followed by injection into 10 equivalents of ice cold water. Concentration and buffer exchange into NMR buffer (25 mM potassium phosphate pH 6.2, 50 mM potassium chloride) was performed using Vivaspin concentrators (MW cutoff 3000 Da, Sartorius).

Commercially obtained chemically synthesized NSR variants with single phosphorothioate groups (Horizon Discovery Ltd.) were deprotected according to the instructions of the manufacturer and folded, concentrated and rebuffered as described previously. The chemically synthesized RNAs have 5′-OH and a 3′-OH groups at their termini, causing the imino proton resonances for the first two base pairs to be slightly shifted compared to the *in vitro* transcribed NSR variants.

### NMR spectroscopy measurements

The NMR sample concentrations of RNA varied between 150 and 460 μM and were titrated with 1 to 1.5 equivalents of commercially available ribostamycin (Merck). The RNA saturation with the ligand was confirmed by the disappearance of imino proton resonances of the free RNA in 1D-^1^H-NMR-spectra. All NMR spectra were collected on 600 MHz and 700 MHz Bruker Avance NMR-spectrometers equipped with 5-mm cryogenic triple resonance TCI-N and TCI-P or quadruple resonance QCI-P probe heads. All NMR experiments were recorded at 10°C in 5% (v/v) D2O / 95% H2O in a buffer containing 25 mM potassium phosphate and 50 mM KCl, pH 6.2. All NMR data were processed using Topspin 4.0.6 (Bruker Biospin). Most of the imino proton resonance assignments of the NSR variants could be transferred from earlier measurements of the wt and mutC14 variants.^30,38^ Remaining imino resonances could be assigned with imino-imino cross-peaks of 2D-^1^H,^1^H-NOESY spectra. 2D-^1^H,^1^H-NOESY, 2D-^1^H,^15^N-HSQC and 2D-H(N)C experiments were recorded using standard pulse sequences.^72^ For detection of NH…O-P and NH…S-P hydrogen bonds, long-range 2D-^1^H,^31^P-HSQC experiments^73,74^ were carried out.

## Results and Discussion

The goal of our work is a comprehensive characterization of phosphorothioate (PT) modification in the context of a folded RNA molecule (Neomycin-sensing riboswitch, NSR) using molecular dynamics simulations, QM/MM and QM computations.

We first present NMR experimental measurements demonstrating the validity of studying PTs in context of the NSR using computational approaches. Then, we describe MD simulations of the phosphorothioate (PT) modification in the NSR and compare them with simulations of unmodified systems. We show how PT influences the structure and dynamics of the NSR and the conserved interactions within its U-turn motif – the HB1 and HB2, the anion-π stacking, and the potassium binding site (Figure 1 and Figure 2). Structural and dynamical difference between the unmodified RNA and the phosphorothioate modified RNAs were mainly observed for the Rp stereoisomers of PTs which are reported in the main text while the Sp stereoisomers are discussed in the Supporting Information. We then verify the simulation performance by extensive QM/MM optimizations and interaction energy QM scans of the full NSR and smaller model systems, respectively. The electronic structure calculations provide insights into the physical nature of PT molecular interactions and explain why its correct description using MM parameters is challenging. We show that MM description can be partially improved by using NBfix. Finally, we present series of advanced NMR measurements and QM/MM computations to investigate the interplay between the PT and U14C substitution.

### PT replacements in the RNA backbone are compatible with the NSR fold

All our initial structures of NSR with PT were obtained by modeling the PT modification into a folded wild-type structure of the NSR (see the Methods). While this is a commonly utilized approach in computational studies,^20^ it assumes that the PT modification at phosphates A16 and A17 does not destabilize or radically alter the RNA fold of the NSR.

To experimentally verify these assumptions, we recorded ^1^H-NMR spectra of the NSR bound to ribostamycin with PTs at either A16 or A17, respectively. The chemically synthesized RNAs contained a 1:1 mixture of the Rp and the Sp stereoisomers (see the Methods). As a result, the measured NMR-spectra featured two sets of signals for atoms in the vicinity of the substitution site. For the system with PT at A17, we see double signals for the imino protons of U13, U14 and U18 (Figure 3B, top) compared to single signals for these nucleotides in the wild-type RNA (Figure 3A). All other imino proton signals belong to atoms further away from the PT site and appear as single signals at positions corresponding closely to the wild-type NSR^75^, indicating that they are not affected by the PT and that its influence is confined to the U-turn loop. The presence of two imino proton signals for U14 indicates that it is engaged in a hydrogen bond in both the Sp and the Rp stereoisomers, indirectly pointing to the presence of the signature HB1 interaction; in case of Rp with the sulphur acting as HB1 acceptor. To directly confirm the presence of HB1 with the A17 PT, we performed a long-range 2D-H,P-correlation experiment where magnetization is transferred through scalar couplings across the hydrogen bond.^73,74^ There, we observe cross peaks (Figure 3B) for the U14 imino resonances from both stereoisomers to the signals of the A17 PT group, thus providing direct experimental confirmation of HB1’s presence in both the A17Sp and A17Rp stereoisomers. Further details of these experiments are reported in the Supporting Information. The NMR measurements of the A16 PT system also confirm that the HB1 bond is present with both stereoisomers (Figure 3C), details of which are discussed in the Supporting Information.

**Figure 3:**
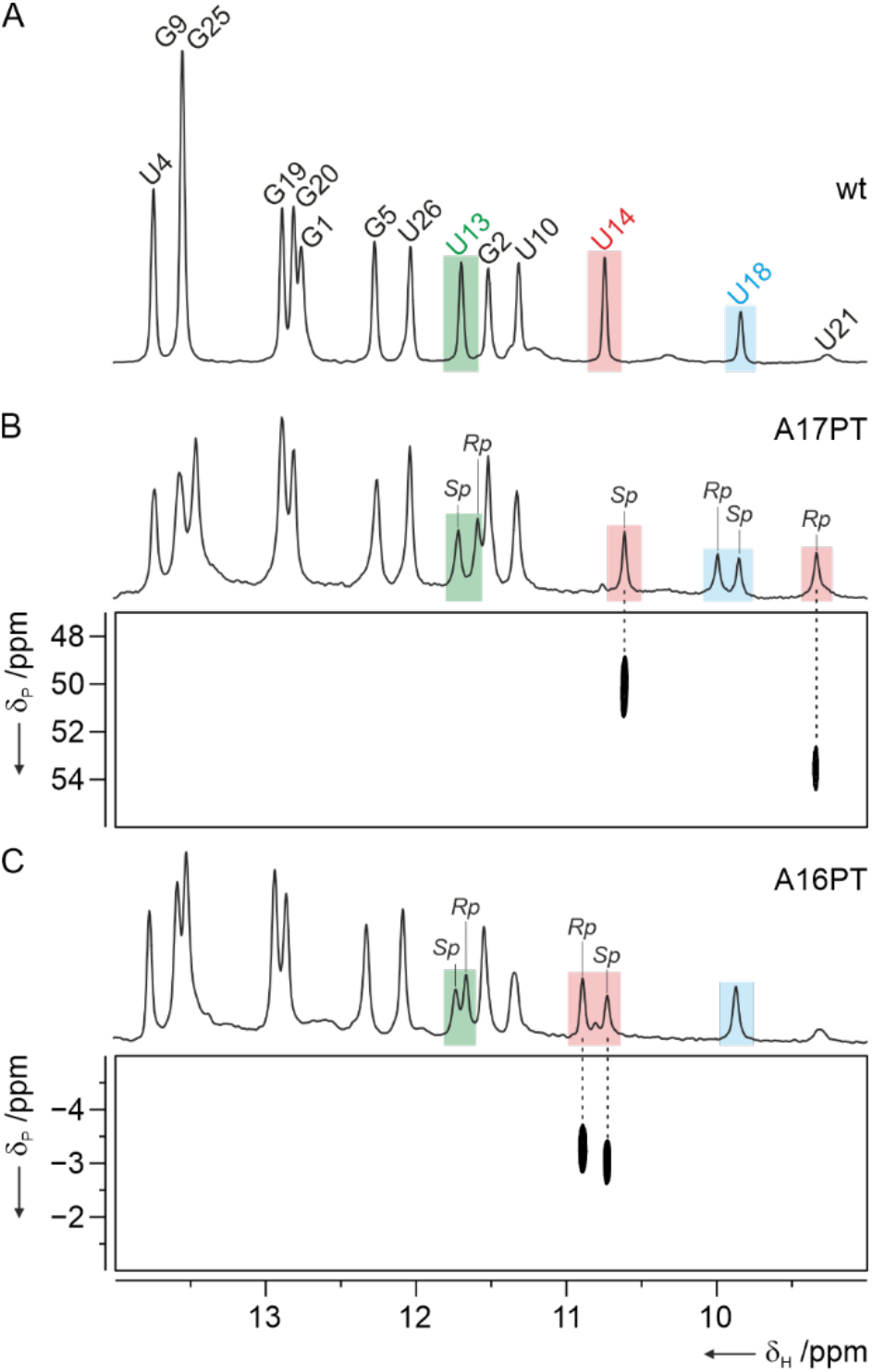
PT replacements at A16 and A17 do not abolish the NSR fold. A) Imino proton spectrum of the wild-type NSR bound to ribostamycin. NMR signal assignments are indicated and the imino protons of U14 involved in the HB1 signature interaction, and U13 and U18 which form the U-turn closing base pair are highlighted in red, green and blue, respectively. B) Imino proton spectrum of the NSR with A17 PT (top). The mixture of Rp and the Sp stereoisomers in the sample produces two sets of signals for the U14, U13 and U18 imino protons. A long-range 2D-H,P-correlation experiment (bottom) reveals scalar couplings across the HB1 interaction in both stereoisomers and confirms that in both stereoisomers, the signature HB1 hydrogen bond is stably formed. C) Imino proton spectrum of the NSR with A16 PT (top). Two sets of signals are observable for the U14 and U13 protons. The A16 PT modification does not affect the U18 imino proton. The 2D-H,P-correlation experiment (bottom) reveals the presence of a stable hydrogen bond corresponding to the HB1 signature interaction in both stereoisomers (bottom).

Overall, the strong resemblance of the imino proton spectra of the Rp and Sp stereoisomers of the A16 PT and A17 PT variants and the wild-type shows that these PTs do not abolish the global fold of the NSR. Furthermore, the H,P-long range correlation experiments directly demonstrate that in all four RNA variants, the signature HB1 hydrogen bond, which is the essential component of the U-turn motif, is present and stable. We suggest this justifies the use of NSR as a suitable computational model to explore the consequences of PT substitutions in RNA. Notably, the observation of the H,P-correlation across the hydrogen bond between the imino proton of U14 and the PT group in the A17 PT variant represents to our knowledge the first direct detection of an NH…S hydrogen bond in RNA by NMR.

### A17 PT tunes the geometry of the HB1 interaction

In the first series of MD simulations, we have utilized the standard AMBER vdW parameters for the sulphur (SP2) atom (Table 3). The PT did not abolish the HB1 and HB2 interactions (Figure 3A) but there were some differences in the geometry compared to the wild-type. Namely, for HB1 in A17Rp systems, we saw a shift in the distribution of its acceptor-donor distances by 0.2-0.3 Å (Figure 4B,C), reflecting the larger vdW radius of sulphur atom compared to oxygen. This minor difference did not seem to have any effect on the NSR overall structure. In A17Rp/C14 and mutC14 systems, the N4 amino group of C14^+^ was commonly observed forming also an C14(N4)/A17(SP2) and C14(N4)/A17(OP2) H-bond, respectively, which we term HB1a H-bond. In simulations, both the HB1 and HB1a can be populated simultaneously, leading to an HB1/HB1a H-bond bifurcation. However, as shown by position density maps (Figure 4D), in majority of simulation frames the acceptor atom was closer to the C14(H3) atom than to C14(H41). We explored the HB1/HB1a H-bond bifurcation further using NMR experiments and QM/MM calculations (see below).

**Figure 4:**
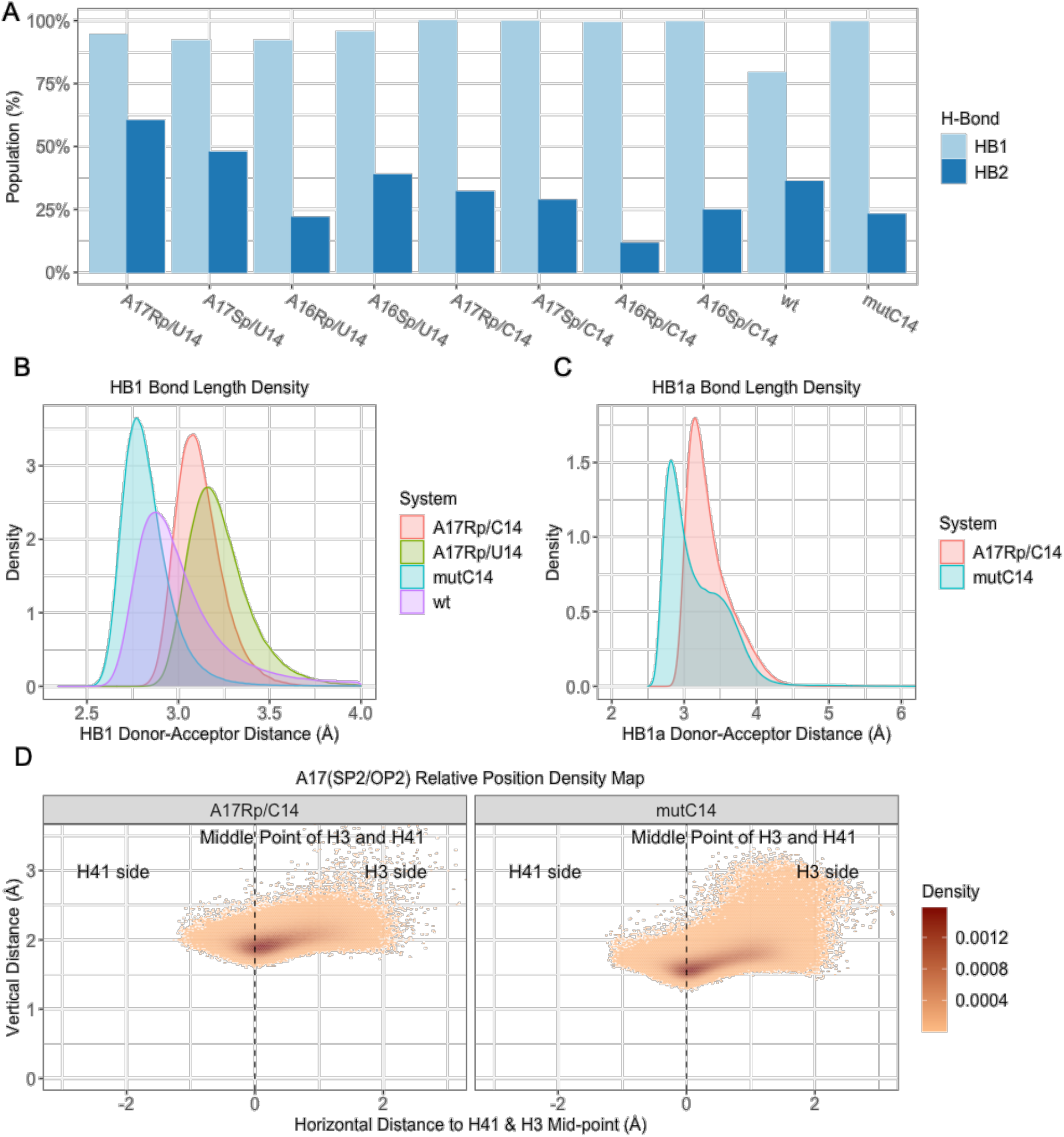
A) Populations of signature H-bonds of the U-turn motif in all simulations (cf. Table 1). B) Density graph of the acceptor-donor distances of HB1. C) Density graph of the acceptor-donor distances of HB1a. D) Projection of the A17(OP2/SP2) atom on the line connecting the C14^+^ H3 and H41 hydrogen atoms. The x and y distance stand for distance of the projection point from the mid-point of the connecting line and separation of the A17(OP2/SP2) atom from the line, respectively.

### *A16 PT increases vertical distance of the anion-*π *interaction*

The anion-π interaction is present in all simulations (Figure 5A). We observed system-specific differences in distance separation (i.e., vertical distance) of the A16(SP2/OP2) atom from the base plane. Namely, the shortest separation was observed in mutC14 simulations and longest in the A16Rp/U14 simulations (Figure 5B). The shorter separation in C14^+^ systems is caused by the protonation and thus the more positive charge of the C14 base.^29^ The larger vdW radius of sulphur then increases the separation distance in A16Rp/U14 and A16Rp/C14 systems compared to the wt and mutC14 systems.

**Figure 5:**
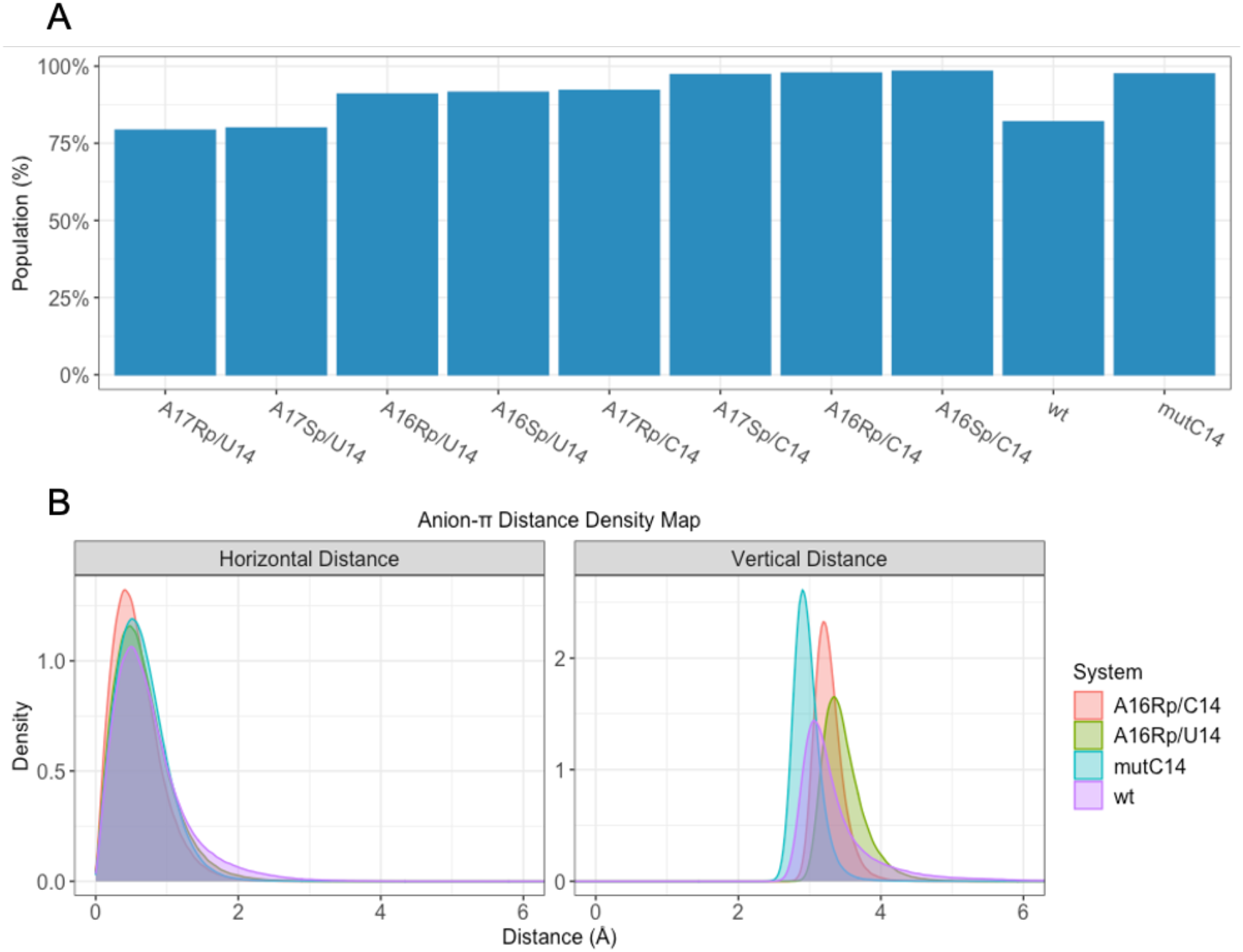
Projection of the A16(OP2/SP2) atom onto the plane of the U14/C14^+^ base. A) Populations of the signature anion-π interaction of the U-turn motif in all simulations (see Methods for definition). B) The horizontal distance (left) and vertical distance (right) are distances between the perpendicularly projected point and the geometric center of the base, and between the A16(OP2/SP2) atom and the base plane, respectively.

### K^+^ binding site near the U14(O4) adjusts its geometry with both A16 and A17 PTs

All simulated systems revealed a major potassium binding site in the proximity of the U14(O4) atom (Figure 2), as visualized by the radial distribution functions (RDFs) in Figure 6, and previously suggested.^29^ The presented RDFs describe the simulation distribution of potassium ions around U13(O4), U14(O4), A16(OP2/SP2), and A17(OP2/SP2) atoms, which together form a highly negatively charged region of the U-turn motif. Compared to the wt, the graphs of A17Rp and A16Rp showed potassium density peaks shifted to greater distances for the A17(SP2) and A16(SP2) atoms, respectively. In case of A16Rp, the ion occupancy for A16(SP2) was also significant compared to the A16(OP2) where it was negligible. Therefore, the PT simulations confirm that the potassium binding site is still present at this location similarly to the wild-type, albeit with slightly altered geometry. Due to the larger vdW radius of the sulphur, the K^+^ is additionally coordinated by A16(SP2) in A16 PT systems. The K^+^ binding site is absent in systems with C14^+^.^29^

**Figure 6:**
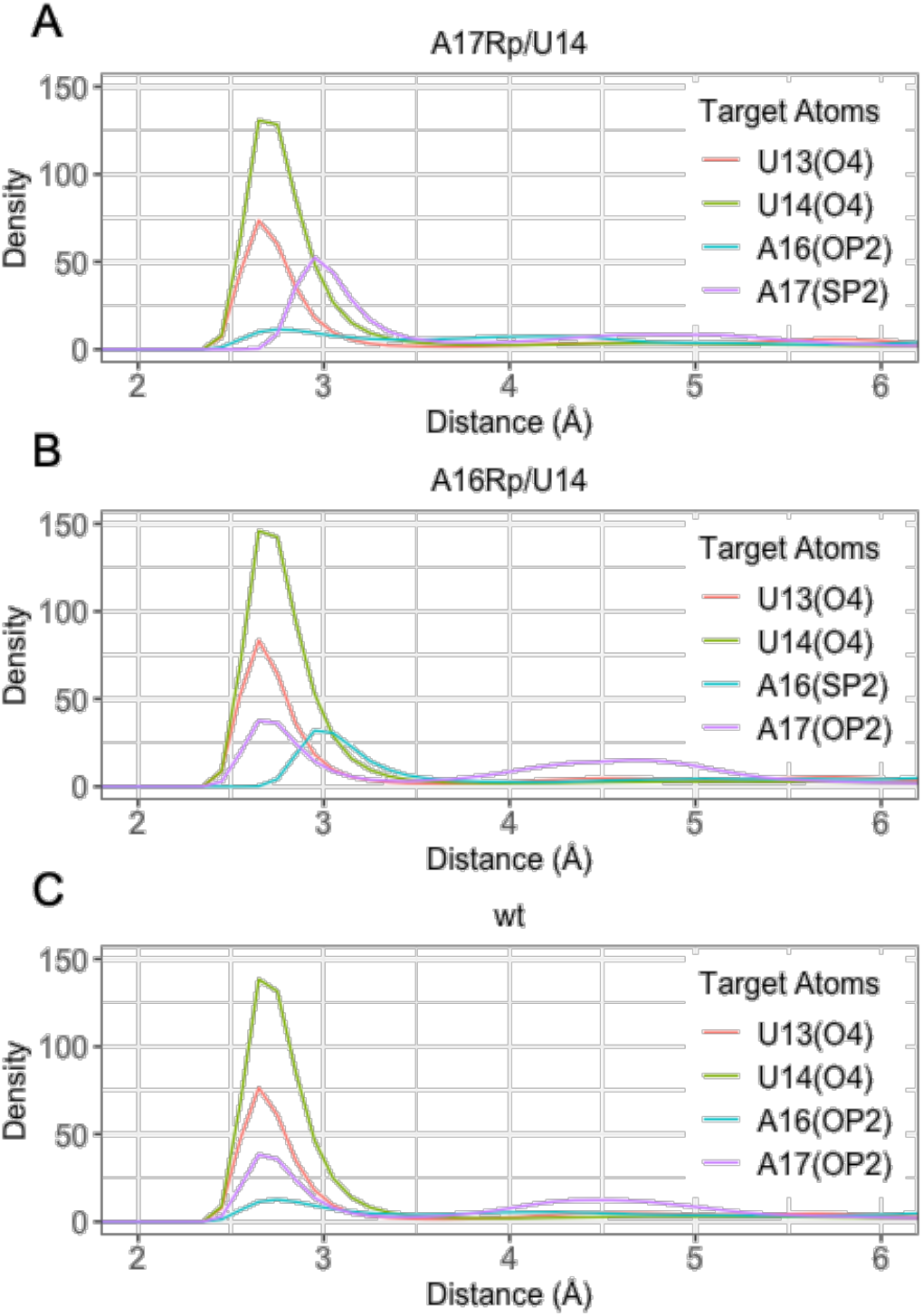
A, B, C) Normalized radial distribution functions of K^+^ ions for the coordinating atoms of the potassium binding site in MD simulations of different systems.

### QM/MM and QM calculations suggest inaccuracies in the MM description of PT when utilizing standard AMBER sulphur parameters

The comparison between QM/MM and MM optimized structures revealed that the acceptor-donor and acceptor-hydrogen distances of HB1 are longer by ca 0.25 Å in QM/MM optimizations of the A17Rp/U14 and A17Rp/C14 systems compared to MM using the standard AMBER parameters (Table S2). The interaction optimum is shifted to longer distances, i.e., the MM sulphur atom is too small. For the remaining systems the difference between the QM/MM and MM geometries is much smaller, i.e., within the expected uncertainty range of such computations.^26^ Thus, the QM/MM optimizations of the full NSR suggested that a vdW parameter for sulphur with a larger radius than the AMBER default would be better for the description of the HB1 interaction in A17Rp systems.

We further carried out QM interaction energy scans of small model systems representing the isolated HB1 interaction as seen in the NSR. We also performed equivalent MM scans utilizing either the original AMBER vdW parameters for the sulphur atom or the thiolate parameters from CHARMM (vdW radii of 2 Å and 2.20 Å, respectively; see Table 3). The calculations revealed that compared to QM in vacuum, the AMBER and thiolate MM parameters under- and overestimated the SP2 – H3 distance in HB1, respectively (Figure 7, S1, Table 4), although the thiolate parameter is closer to reproducing the optimal QM distance. As expected, the QM distance scans showed that the optimal H-bonding distances are shorter for the OP2 – H3 than for the SP2 – H3 (Figure 7, Table 4). The same qualitative difference is observed in MM scans, though the MM (AMBER) description exaggerates the short-range repulsion, as common when using Lennard-Jones potentials (Figure S2).^76^ This contributes to the considerably deeper QM curves compared to MM, which is most visible for the entire range of OP2 – H3 gas phase scans, where polarization also plays a significant role. Taken together, the standard vdW AMBER parameters for sulphur fail to capture the relative differences between sulphur and oxygen atoms in the context of the HB1 interaction. Using the thiolate vdW parameters for sulphur could mitigate this problem and improve simulation description of HB1.

**Table 4:** Summary of the QM and MM HB1 scans of DMTP/DMP – MU distance.

| System | Calculation method | Optimum distance (Å)<sup>a</sup> | Minimum energy (kcal/mol) |
| --- | --- | --- | --- |
| **DMTP – MU** | MM AMBER / vacuum | 2.11 | -3.19 |
|  | MM thiolate / vacuum | 2.38 | -1.96 |
|  | MM AMBER / GBSA | 2.19 | -1.92 |
|  | MM thiolate / GBSA | 2.52 | -1.51 |
|  | QM / vacuum | 2.27 | -6.48 |
|  | QM / COSMO | 2.39 | -2.58 |
| **DMP – MU** | MM AMBER / vacuum | 1.76 | -6.59 |
|  | MM AMBER / GBSA | 1.82 | -3.54 |
|  | QM / vacuum | 1.59 | -15.24 |
|  | QM / COSMO | 1.70 | -5.80 |

<sup>a</sup>The distance between sulphur (or oxygen in DMP) and U14(H3) associated with the lowest interaction energy.

**Figure 7:**
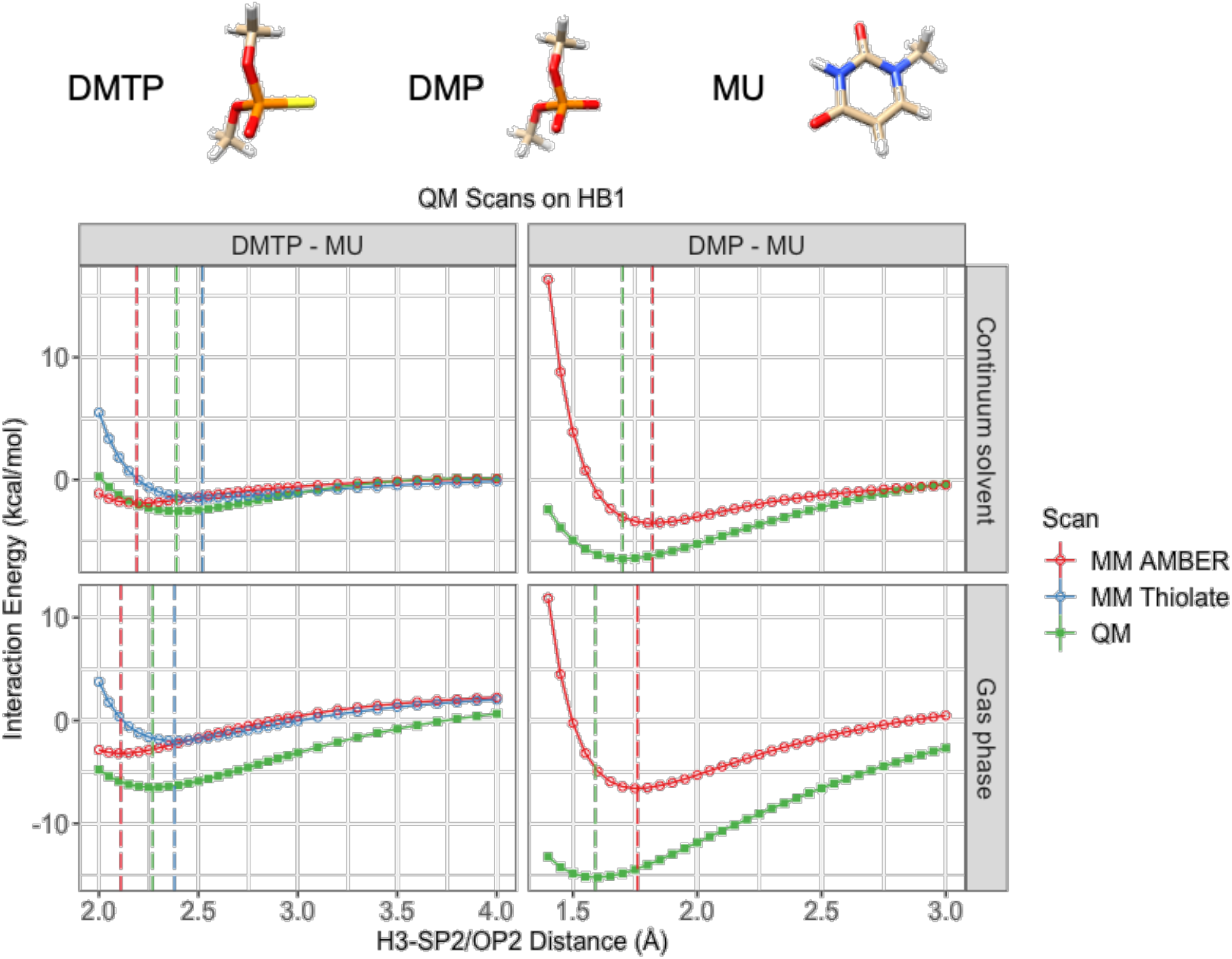
The structures of DMTP, DMP and MU are shown on the top. Interaction energy distance scans (QM PBEh-3c vs. MM) of the HB1 hydrogen bond in DMTP-MU and DMP-MU systems. The vertical dashed lines indicate the position of the energy minimum of each scan. For the complete range of the scans, see the Supporting Information Figure S1. See also Figure S2 for comparison with other QM methods.

It should be noted that obtaining a perfect match between the QM and MM interaction energy curves of the HB1 is not automatically the recipe for a perfect simulation as the simulation behavior will be determined by the balance of all interactions, including solvation. It is therefore necessary to perform simulations of the PT systems with sulphur described by the thiolate parameter (see below) to evaluate its performance. Nevertheless, the failure to capture relative differences between the sulphur and oxygen atoms by the default MM parameters of sulphur is evident from the QM calculations. We also note that the PBEh-3c method agrees with higher-quality QM methods except for a slight overestimation of the absolute interaction energy (Figure S2).

The continuum-solvent calculations are generally known to be sensitive to the used parameters,^20^ making quantitative interpretation of the differences uncertain. So, the results should be considered only as qualitative, to demonstrate over-estimation of the short-range repulsion by the Lennard-Jones term of the force field. For this purpose the accuracy should be sufficient.

A noticable feature of the data in Figure 7 and Table 4 is that the calculations show a much larger difference between the gas phase and continuum solvent curves for the QM calculations compared to the MM computations. Although some discrepancy between the COSMO and GBSA approaches (and their specific parameters) cannot be ruled out, we suggest that the main reason for the difference is the fact that our model systems of the thiophosphate/phosphate interaction with methyluracil are non-neutral, with the overall system charge of −1. This causes significant induction (polarization) effects in QM gas phase calculations while this contribution is zeroed by definition in the pair additive MM. While the QM calculation is physically correct per se, the small-model with unshielded −1 electrostatic interaction does not correspond to an environment in a complete biomolecule.

Considering the potential use of such small-model data in direct parametrization of force fields, the COSMO and GBSA data could be used for some calibration of a force field, as the above-discussed difference originates in the gas phase calculations. Direct utilization of the gas phase calculations for parametrization of a pair additive potential for explicit-solvent simulations is not advised. The data could be relevant in development of polarizable force fields where the parameters would be flexible enough to include the full range of chemical environments, including the gas phase. In case of calibrating pair additive potential, we suggest it would be ideal to use the continuum solvent data in a manner that has been utilized in parametrization of the OL series of nucleic acids force fields, which avoids double counting of the solvation effects.^39,77,78^

### SAPT reveals the distinct natures of the SP2 – H3 and OP2 – H3 HB1 interactions in NSR

To obtain further insights into the physics of the HB1 H-bond, we executed SAPT calculations. The minimum point of the interaction energy of the SP2 – H3 and OP2 – H3 H-bonds returned by the SAPT method is consistent with the PBEh-3c vacuum results (Figure 8, Table S4). The energy decomposition indicates that the induction energy is the main binding contributor to the SP2 – H3 H-bond, while for the OP2 – H3 bond, it is electrostatic energy. The dispersion energy with sulphur is also more significant than with oxygen (Figure 8 and Table S4). These trends are observed throughout the scanning range and reflect the greater charge-density extent and polarizability of sulphur compared to oxygen.

**Figure 8:**
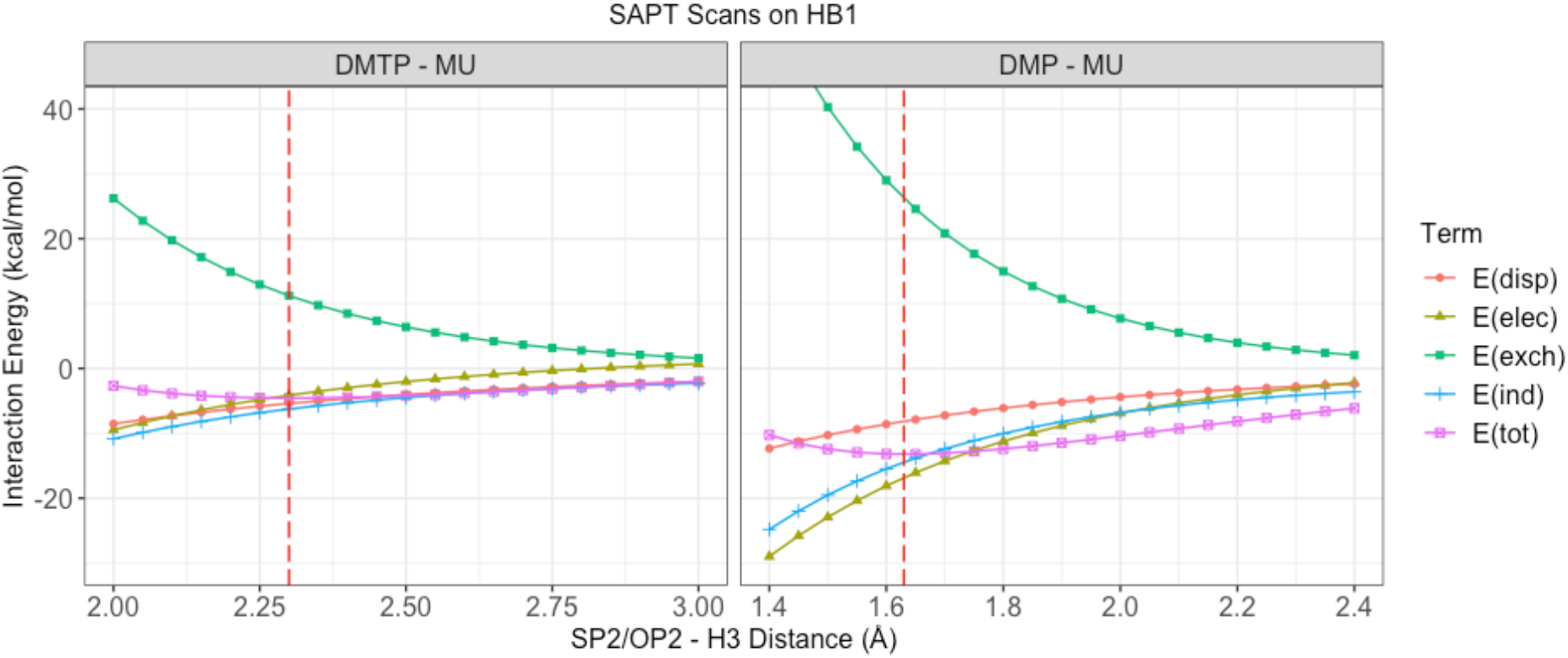
QM scans of the SP2 – H3 – N3 and OP2 – H3 – N3 H-bonds (HB1), respectively, using the SAPT method. The red dash line notes the minimum of the total energy.

### PT relaxes directionality requirements for the HB1 signature H-bond

The SAPT calculations of the HB1 model systems demonstrated the increased polarizability of the sulphur compared to oxygen (Figure 8). Based on this, we hypothesized that the directionality requirements of the H-bonding interaction could be less strict with sulphur as acceptor since the donor polarizes it more readily. To explore this idea, we performed QM distance scans on the DMTP/DMP – MU model in two scanning directions perpendicular to the HB1 bond vector (Figure 9). For each distance point, the HB1 distance was optimized along the HB1 vector (defined by SP2/OP2 – H3) by translating MU in relation to DMTP/DMP. Most interestingly, we found that the geometry associated with the optimal interaction energy for the DMTP – MU is shifted outside of the direct (180°) angle of the HB1 by at least 0.6 Å in both directions. In contrast, the DMP – MU model showed the optimal interaction energy for an HB1 geometry close to the direct angle (Figure 9). The interaction energy gradient was also steeper with DMP in both scanning directions.

**Figure 9:**
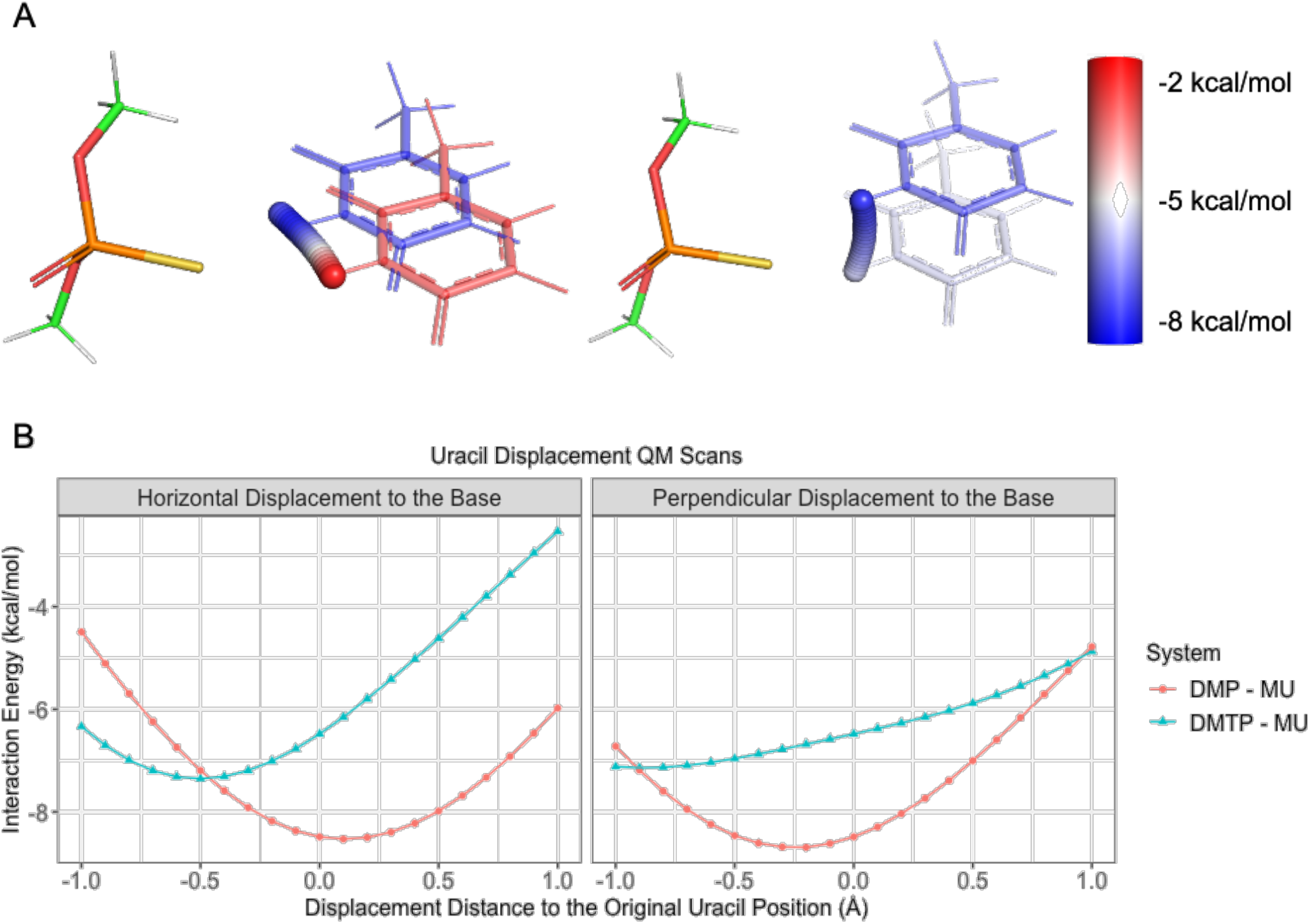
A) Interaction energy scans between DMTP and MU in two directions (left: displacement horizontal to the base, right: displacement perpendicular to the base); the inter-monomer distance is relaxed in each point (see the text). The interaction energies are visualized by the colors on the H3 spheres for each uracil displacement. From red to blue, the interaction energy decreases. The rest of the base is displayed only for positions associated with maximum and minimum interaction energy, respectively. B) Graphs of QM interaction energy scans of MU displacement against DMTP/DMP.

In conclusion, these data confirm that the HB1 interaction with DMTP is inherently less directional and has a lower potential energy penalty for HB1 angle changes than with DMP. We suggest that in full biomolecular systems, such interaction could better tolerate thermal fluctuations even though the interaction energy is intrinsically weaker with sulphur. However, how such an effect would change the thermodynamic stability of the system would obviously also depend on its full structural context, the balance of solute-solvent interactions, and interactions in the unfolded state. However, the sulphur acceptor seems to better tolerate reduced directionality of H-bonding and may also support hydrogen bond bifurcation more readily than oxygen (see below).

### Thiolate parameters improve the agreement between MM and QM also for the PT interaction with water

Our distance scans of the HB1 model tentatively suggested that the thiolate parameters in MM calculations are better for reproducing the QM results. We have subsequently calculated QM and MM distance scans between the DMTP/DMP and a water molecule to examine the MM parameters performance in more chemical environments (Figure S3). This is important as in addition to forming HB1 interaction in the NSR, the sulphur atom is also simultaneously interacting with the solvent. These calculations showed that the standard AMBER sulphur parameters give a too short H-bonding distance also for the SP2 – water interaction (Table S5, Figure S4) and that the use of thiolate parameters provides a much better fit to the QM data. Based on this data we suggest that the thiolate parameters are a superior choice in MD simulations, for both describing the signature HB1 interaction, as well as the water interaction with the sulphur atom of PT. The QM and MM scans of the OP2 – water interaction distance indicated an overall satisfactory performance of the force field (Table S5, Figure S4).

### The QM distance scans of the anion-π interaction reveal opposite requirements for the sulphur Lennard-Jones radius in MM than for the HB1 interaction

The anion-π interaction QM distance scans were done on small models extracted from the QM/MM optimized A17Rp/U14 and A16Rp/U14 structures. DMTP/DMP was separated from MU in the direction perpendicular to the base plane to obtain the interaction energy profiles. For both systems, the QM indicates shorter optimal SP2/OP2 – base distances than the MM (Figure 10), reflecting a rather considerable overestimation of the short-range repulsion by the Lennard-Jones potential. As a result, unlike for the HB1 interaction (see above) the use of thiolate parameters with the bigger vdW radius caused the anion-π distance in MM to further deviate from the QM value. It suggests that a simultaneous accurate description of both the HB1 and the anion-π interaction through a single vdW radius of sulphur might not be possible.

**Figure 10:**
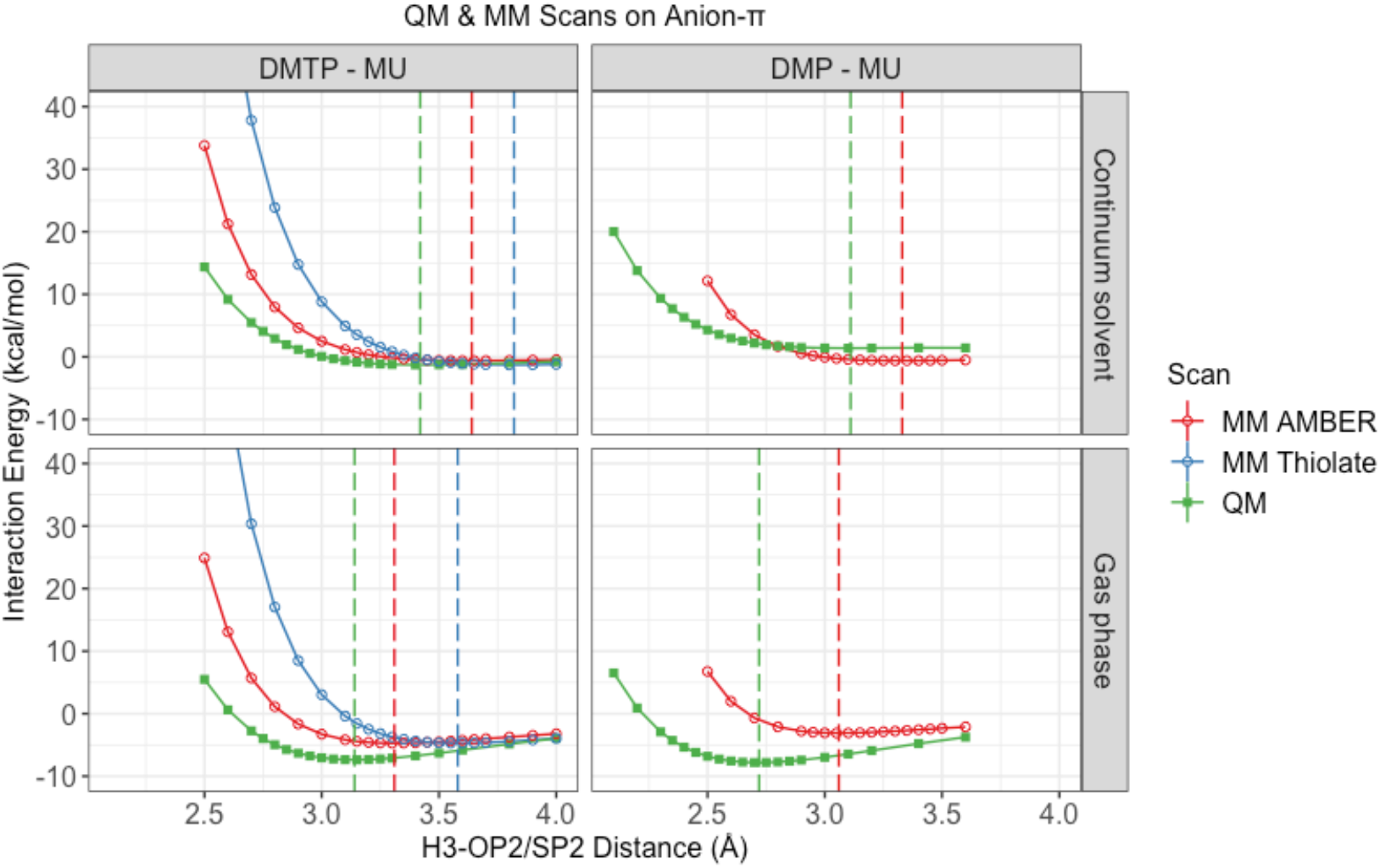
The anion-π distance scans for the DMTP/DMP – MU model system in implicit solvent (top) and vacuum (bottom). The vertical dashed lines indicate the position of the energy minimum of each scan.

We have also carried out SAPT energy decomposition of the anion-π interaction model systems. The electrostatics contribute to the SP2 – base anion-π interaction more than for the OP2 – base interaction (Figure 11, Table S6). However, the OP2 – base interaction is more stabilized by induction, suggesting that oxygen is able to considerably better polarize the base than the sulphur. These energy contributions show virtually opposite trends compared to the HB1 interaction (Figure 8), providing an explanation why a single vdW parameter cannot ideally describe sulphur in both chemical environments. It should be noted that gas phase interaction energy decomposition of non-neutral systems might exaggerate the role of electrostatics and induction components compared to biochemically relevant environments.^79^ To the best of our knowledge no SAPT theory accounting for solvation effects is available to include, e.g., the role of polar solvent screening. The presented results are correct per se but assume no solvent screening. Neutral systems also experience certain damping effects upon inclusion of solvent screening, but to a significantly smaller extent, making their SAPT results easier to interpret and extrapolate. However, the basic message of our SAPT calculation should be correct.

**Figure 11:**
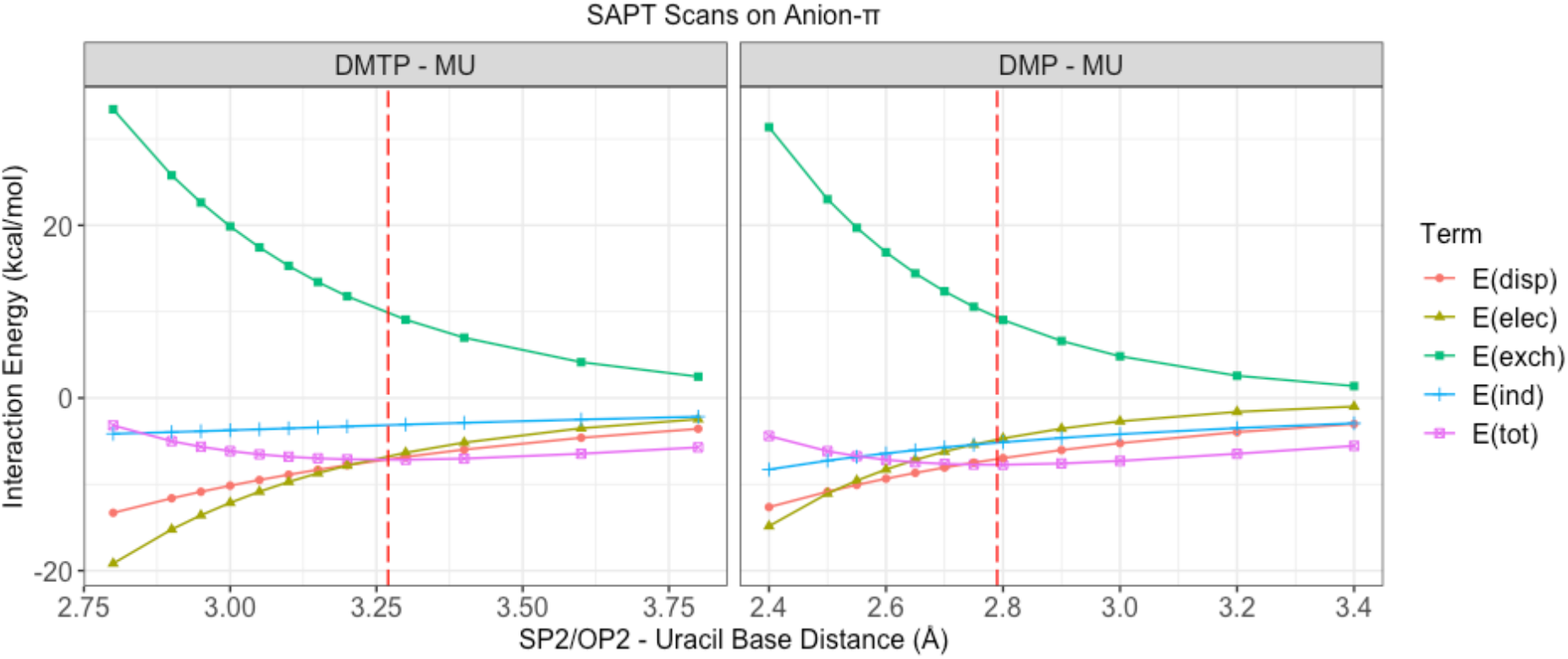
Interaction energy scan by SAPT for the anion-π interaction of A16(SP2) (left)/A16(OP2) (right) – U14. The vertical red dashed line indicates the minimum of the total energy.

In conclusion, for the anion-π interaction, except of the short-distance repulsion, high-quality QM data agree quite well with the MM scan using AMBER parameters (Figure S5). Therefore, unlike with the HB1, the use of the AMBER sulphur parameters for the description of the anion-π interaction can be considered appropriate while the thiolate parameters do not seem suitable. Over-estimation of the short-range repulsion is common for Lennard-Jones potentials^76^ and appears to be larger for the oxygen atom. Potential impact of this effect on description of anion-π interactions in MM calculations will be addressed in future studies.

### Adjusting the sulphur vdW parameters for PT in MD simulations by thiolate parameters and NBfix

The QM/MM and QM calculations suggested two incompatible requirements for the ideal vdW parameters of the sulphur atom in MM, with the HB1 interaction requiring a bigger (see above and Table 4, Figure 7 Figure *8*) and the anion-π interaction a smaller Lennard-Jones radius in order to match the QM data. These results nicely illustrate the complicated chemical features of sulphur and the general difficulties in optimizing a single force-field parameter to perform satisfactorily in multiple chemical environments. Based on the above QM/MM and QM calculations, we tried to propose a solution to the conundrum of sulphur vdW parameters which would be ideal for simulations of all PT-containing systems. As summarized in Table 3, in our MD simulations we tried global implementations of the AMBER and thiolate parameters, respectively, as well as NBfix for specific pairwise interactions of the sulphur. With NBfix, the thiolate parameters were implemented for nonbonded interactions between the sulphur and N3 and H3 atoms of uracil as well as the solvent molecules while the AMBER parameters were used for all the other nonbonded interactions formed by sulphur. We note that it is necessary to use the same vdW parameter for the interaction with all solvent particles (i.e., water and ions) to avoid serious imbalances. The A17Rp/U14 simulation with the AMBER sulphur vdW parameter has the shortest HB1 length, while the A17Rp/U14 simulations with either globally or the NBfix-implemented thiolate parameters have approximately 0.3 Å longer HB1 length (Figure 12A). Importantly, the comparison between globally and NBfix-implemented thiolate parameters further shows that the NBfix implementation improves HB1 without worsening the anion-π interaction in the A16Rp/U14 system (Figure 12B). The NBfix approach is thus an appropriate method to handle the opposite demands for the vdW radius of the sulphur atom in both chemical environments. Further details of the MD simulations utilizing the thiolate sulphur parameters, either globally implemented or by NBfix, are presented in the Supporting Information.

**Figure 12:**
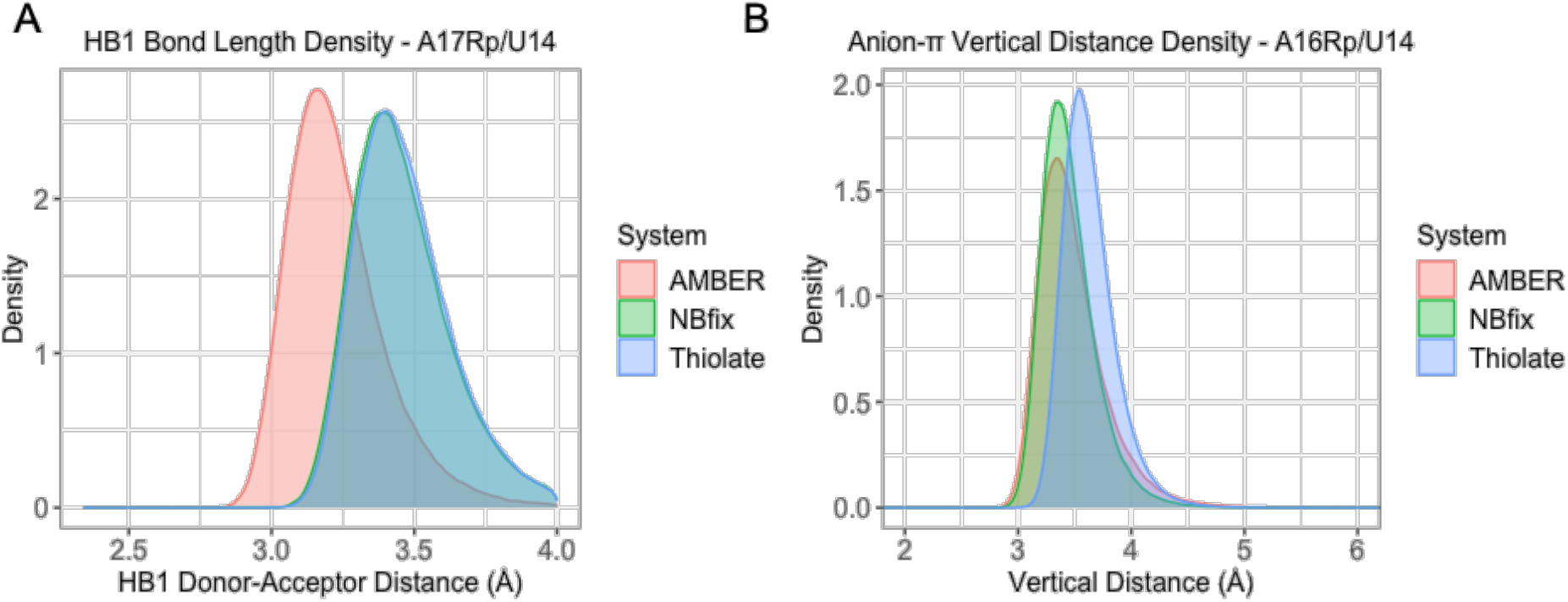
Comparison of HB1 and anion-π interactions in simulations with different vdW parameters for sulphur. A) HB1 hydrogen-acceptor distance density in the A17Rp/U14 simulations. The NBfix and thiolate curves are overlapped. B) The anion-π vertical distance density in the A16Rp/U14 simulations. For additional discussion of the differences, see the Supporting Information.

### NMR measurements of the C14 NSR show the A17Rp PT can influence the pK_a_ elevation of C14

The U14 in the NSR U-turn loop can be replaced by a protonated C14 without altering its structure (mutC14 system; see Table 1). The C14 protonation is stable and the H3 proton clearly detectable by NMR even at pH values > 8.3, which is remarkable considering the pK_a_ of free cytosine nucleotides (~ 4.5) (for more details, see the Introduction and Refs.^29,38^). This local elevation of the C14 pK_a_ in NSR entirely depends on the chemical environment of the C14(N3) atom in fully folded NSR and could therefore be significantly altered by the A17 as well as A16 PTs, which directly impact the signature HB1 and anion-π interactions, respectively. This is of particular concern in the case of the two Rp stereoisomers where the sulphur atom is in direct molecular contact with C14 base.

In contrast to the PT-modified NSR with U14, we could detect only a single signal by the NMR for the imino proton of the protonated C14 in both the A16 and the A17 PT-modified RNAs. We also observe a single correlation in the respective long-range H,P-correlation spectra (Figure S6B). For both the A16 and A17 PTs, we assign these signals to the Sp stereoisomers. This could potentially suggest that in the Rp stereoisomers, the C14 is either no longer protonated or that its imino proton is in fast or intermediate exchange with the bulk water and its signal broadened beyond detection. In both cases, the U-turn loop would not be properly folded under such circumstances. Still, comparison of the imino proton spectra of the mutC14 NSR (Figure S6A) and chemically synthesized A17 PT and A16 PT RNAs showed that the presence of the modification leads to splitting of the U13 imino proton signal for both RNAs and of the U18 imino proton signal for the A17 PT RNA (Figure S6B) in the ligand-bound state. This is analogous to what was observed upon introduction of these PTs into the U14 NSR and confirms that at the very least the U13/U18 base pair is not disturbed by any of the PT stereoisomers, even in the presence of the C14 mutation.

Therefore, to gain clear insight into the actual folding state of the U-turn loop, we examined its ^31^P-NMR-spectra. The deviation from A-form helical torsion angles for nucleotides U15, A16, A17 and U18 associated with the U-turn loop is reflected in unusual upfield or downfield chemical shifts from the bulk of A-form helical ^31^P-resonances in the range between −0.5 and −2.0 ppm in the unmodified RNA.^38,80^ The ^31^P-NMR spectrum of the A16 PT NSR with C14 (Figure S7A) showed two signals for the U15 and U18 phosphate groups which are shifted downfield as well as two signals for the A17 phosphate shifted upfield compared to the bulk of the ^31^P signals. Furthermore, two sharp signals are observed for the A16 PT group with ^31^P chemical shifts > 50 ppm. This confirms that the A16 PT backbone torsion angles and conformation closely resembles the mutC14 and wt RNA and that the U-turn loop is folded in both A16Sp and A16Rp even though the C14 protonation is not directly observable for the Rp stereoisomer.

The situation is more complex for the A17 PT where we observe only a single signal for the phosphate groups of U15, A16 and U18 as well as for the A17 PT group (Figure S7B). Taken together with the absence of a signal for the imino group of the protonated C14 in the A17Rp, this suggests that under the conditions of our measurement (pH 6.2), the C14 is not stably protonated when the sulfur atom acts as the HB1 acceptor and the U-turn loop is unstable in the A17Rp. In particular, the complete absence of a signal for the A17 phosphorothioate group suggests that there is conformational exchange in the loop on the intermediate exchange time scale for NMR that leads to the loss of this signal due to excessive exchange-induced line broadening.

We subsequently prepared C14 NSR containing only the Rp stereoisomer of A17 PT (see Supporting Information) and performed heteronuclear experiments using the ^15^N and ^13^C-nuclei of the cytosine nucleotides^81^ which can unambiguously establish the protonation state of C14 by observing the ^13^C chemical shift of its C4 carbon in a 2D-H(N)C correlation experiment even when the H3 proton signal is broadened beyond the detection limit.^38^ These experiments confirmed that the C14 is indeed unprotonated in the A17Rp/C14 system at pH 6.2, which was the standard buffer value for our measurements. We subsequently tried lowering the pH to 5.5 and observed that at this pH value, the C14 is once again fully protonated. Similarly, ^31^P-NMR spectra of this RNA taken at pH-values of 6.2, 6.0 and 5.5 reveal the appearance of a signal for the A17 phosphorothioate at lower pH as well as the gradual sharpening of the signals for U15, A16 and U18, in agreement with stable loop folding at lower pH (Figure S8). In conclusion, the A17Rp PT, where the sulfur atom acts as HB1 acceptor, still increases the pK_a_ for the C14 protonation compared to free cytosine, albeit to a significantly lesser degree than the oxygen HB1 acceptor in the mutC14 or A17Sp/C14 systems. A possible cause of this is a putative bifurcation of the HB1 interaction between the N3 imino and N4 amino groups in the A17Rp/C14 system which was strongly suggested by the ^15^N-HSQC spectra of the amino groups (Figure S9C and Supporting Information). We study the idea of the HB1 bifurcation further in our QM/MM calculations.

### QM/MM confirms that the HB1 bifurcation in C14^+^ systems could be realistic with the PT on A17

In our earlier study, we have investigated the potential existence of the HB1/HB1a bifurcated interaction seen in MD (Figure 4D) in the mutC14 system by QM/MM and concluded that the minimum of potential energy is associated with the C14(N3) atom acting as the sole donor. The NMR measurements could not detect HB1/HB1a bifurcation in mutC14 either, suggesting that the force field over-stabilizes bifurcation due to lack of true H-bond directionality in MM.^29^ However, the larger size of the sulphur atom and its different electronic structure could easily change the balance of the interactions in favor of the bifurcation for the A17Rp variant, as was also indicated by the NMR measurements presented here (see above). Therefore, we carried out QM/MM optimizations of the simulation snapshots of individual NSR variants containing the bifurcated HB1/HB1a interaction.

Of all the variants, only the A17Rp/C14 and A17Sp/C14 systems maintain the bifurcated arrangement after optimizations in both QM/MM and MM (Table S3). The result for the A17Rp/C14 system shows that the birfurcated HB1/HB1a with sulphur as the H-bond acceptor is at least a local minimum on the potential energy surface. It could suggest that sulphur, in general, has a larger potential to form bifurcated H-bonds than the oxygen acceptor. This result is supported by our QM calculations on small models which showed that the sulphur has lower requirements for H-bond directionality than the oxygen (see above and Figure 9) as well as by the NMR measurements which detected involvement of the C14 amino group in the HB1 interaction for the A17Rp/C14 system (Figure S9C).

The result of the A17Sp/C14 is more difficult to assess as we do not have a way to experimentally prepare an RNA which would contain solely the Sp stereoisomer of PT. Therefore, to verify our computational result we further performed re-optimization starting from the A17Sp/C14 optimized structure in which we replaced the sulphur with oxygen. In this optimization, the bifurcation was mitigated, suggesting that the bifurcation of HB1 in A17Sp/C14 might be related to some electronic structure differences of the thiophosphate, even though the sulphur is not acting as the acceptor atom (Table S3). However, such result should be interpreted within the sampling limits of QM/MM optimizations.^26^

For the systems besides A17Rp/C14 and A17Sp/C14, the HB1a interaction disappears in QM/MM in favor of a pure HB1 interaction while the bifurcation is maintained in MM. For the unmodified mutC14 system, this is consistent with the earlier study.^29^ In summary, all the experimental and computed data suggest the MM exaggerates the tendency to form bifurcated H-bonds by C14^+^, though with the A17 PT modification, some degree of bifurcation could be expected in real molecules.

## Concluding remarks

We used MD, QM/MM, and QM calculations and NMR spectroscopy to study phosphorothioation (PT) in the NSR, the smallest biologically functional and also structurally well-defined riboswitch, providing new information on the effects of this chemical modification on RNA structure.

We first showed by NMR that the PTs do not alter the overall fold of the NSR. We then used computations to reveal that the conserved interactions of the NSR’s U-turn motif are maintained but their geometries are modulated with the PT, including the signature H-bonds, the anion-π interaction, and the potassium binding site. Although the PT on the A17 phosphate group maintains the signature H-bonds, the distance between the donor and acceptor is extended compared with the wild-type. This may be compensated for by less strict H-bond directionality requirements of the sulphur acceptor which is also reflected in bifurcation of the HB1 interaction in C14^+^ NSR system when PT is the acceptor, as suggested by both QM/MM calculations and NMR measurements. In the A16Rp/U14 and A16Rp/C14 systems where PT occurs at the A16 phosphate, the anion-π interaction remains stable. However, the sulphur is farther away from the U/C14^+^ base compared to anion-π interaction involving oxygen atom.

The MD results would suggest that the effect of PT is related to the larger vdW radius of the sulphur and the associated alterations of U-turn geometry. However, the MM approximation cannot directly reproduce effects related to the different electronic structures of the phosphate and thiophosphate.^20,76^ These differences cannot be always accurately described using simple approximations like atomic point charges and Lennard-Jones potentials. To remedy this drawback, we have performed QM/MM optimizations on all the simulated systems, comparing them with MM optimized results. The QM/MM results showed a longer distance for the HB1 interaction when the sulphur is involved in the H-bond compared to MM. It suggested that a larger vdW radius of the sulphur atom might improve the HB1 description in MD simulations of NSR with PT. This is also well supported by the QM distance scans of small model systems which suggested increasing the vdW radius of the sulphur when it is involved in the HB1 interaction. However, the opposite observation was made for the anion-π interaction where the sulphur vdW radius is already rather large even with the default AMBER force field. In other words, the QM calculations reveal opposing demands on the radius of the sulphur vdW parameters, depending on its chemical environment. This nicely illustrates the inherent problem in force-field descriptions where it might not be possible to always derive a generally satisfactory and universally applicable parameter. Therefore, we suggest implementing the larger sulphur MM vdW parameters in form on an NBfix, modulating only the HB1 and interactions with the solvent, while not altering the systems where the PT is involved in the anion-π interaction. The MD simulations with the sulphur vdW parameters implemented via NBfix describe the HB1, anion-π, and solvent interactions in a manner more consistent with the QM results. We suggest that, in principle, polarizable force fields could be better equipped for dealing with the complex chemical nature of sulphur and the interactions formed by the PTs in MD simulations. The QM/MM and QM data presented here could serve as benchmark for the development of such force fields.^82–84^ We note that anisotropy of sulphur interactions, in comparison with MM descriptions, has also been noted in QM studies of stacking and H-bonding properties of thioguanine and thiouracils.^85^

It should be noted that our μs-scale simulations of the folded state of PT-containing NSR were ultimately satisfactory even with the original sulphur parameters and that the simulations with the alternative sulphur parameters produced highly similar results. We suggest that this reflects the fact that NSR is a well-behaving system in MD simulations, tolerant of minor force-field inaccuracies when simulating just the folded state. However, the imbalance of the standard sulphur parameters could cause misinterpretations when modeling PT modifications in other RNA and DNA contexts where more accurate interaction geometries are necessary or the systems are more flexible, such as in studies of folding or catalysis.

## Supporting Information

The following files are available free of charge https://pubs.acs.org. Details about MD protocol, NMR experiments, MD simulations and QM calculations of systems with Sp PT configuration, additional details about dimethyl-thiophosphate – water interaction energy scans, computational results of the optimized HB2 description with HBfix protocol, details of the interaction energy QM and MM scans, additional MD simulations with different sulphur parameters, incomplete NBfix and its effects on solute-solvent interactions, supporting tables and figures (PDF). Video showing the optimized dimethyl-thiophosphate-water orientations with a series of intermolecular distances given by QM and MM optimization.

## ACKNOWLEDGMENT

This work was supported by the Czech Science Foundation grant 20-16554S (J.S. and M.K.), by H2020-MSCA-ITN project 765266 (LightDyNAmics) by European Commission (Z.Z.), and by project SYMBIT reg. number CZ.02.1.01/0.0/0.0/15_003/0000477 by the ERDF (H.K, J.S). All NMR experiments were conducted at the Center for Biomolecular Magnetic Resonance (BMRZ) at Goethe-University Frankfurt which is supported by the State of Hesse. The acquisition of the quadruple resonance QCI-P cryoprobe required for ^31^P-measurements was supported by the Deutsche Forschungsgemeinschaft (DFG) through Grant INST 161/816-1 FUGG.

