## Supplementary Information for "Phosphorothioate Substitutions in RNA Structure Studied by Molecular Dynamics Simulations, QM/MM Calculations and NMR Experiments"

***Supplementary Text***

*The protocol of the MD simulation.*

The MD simulations were run in AMBER 18.[1] Before the production stage of the simulations, each system was processed by a series of minimizations and equilibrations. First, the minimizations of the systems were done for all atoms with 25 kcal/mol energy positional restraint on the solute. After heating up the systems from 0 K to 300 K with a 25 kcal/mol energy positional restraint on the solute, the subsequent minimization and equilibration cycles were implemented alternatively for five rounds. During this process, the positional restraints on the solute were gradually reduced from 5 kcal/mol to 1 kcal/mol, using a 1 kcal/mol increment. The positional restraint for the final equilibration was 0.5 kcal/mol. For each minimization, we ran 500 steps of steepest descent followed by 500 steps of conjugated gradient minimization. The equilibration steps lasted 50 picoseconds each.

*Details of NMR experiments with the A17 PT NSR.*

For the 2D-H,P-correlation experiments with the A17 PT, we note that the $^{31}$P chemical shifts of the thiophosphate groups in both stereoisomers can be easily identified due to their chemical shifts of ~ 50 ppm.[2] In contrast, the unsubstituted phosphate groups in RNA have chemical shifts in the range of ~2 ppm to -5 ppm. The two imino proton signals of U14 in the A17 PT NSR have a chemical shift difference of ~ 1.4 ppm (9.3 ppm and 11.7 ppm compared to 11.7 ppm in the wild-type) most likely due to the different chemical nature of the HB1 acceptor atom. To confirm which signal belongs to which stereoisomer, we compared the imino proton spectrum of the chemically synthesized RNA containing both stereoisomers with the spectrum of an RNA produced enzymatically by *in vitro* transcription with T7-RNA polymerase in the presence of α-thio-ATP. T7-RNA polymerase produces NSR RNA with three phosphorothioate substitutions in its backbone at A24, A16 and A17 while at the same time, it only produces the Rp-stereoisomer.[2] The A24 is located in the A-form helical stem of the NSR, far away from the ligand binding site and the U-turn loop, and we assume that the PT substitution at this position has no influence on the structure and NMR-spectroscopic properties of the U-turn loop. The imino proton spectrum of this RNA yielded a signal for U14 at ~9.2 ppm but not at 11.7 ppm (Figure S10A). This confirms that the imino proton signal at 9.3 ppm in the A17 PT variant belongs to the Rp stereoisomer whereas the U14 imino proton signal at 11.7 ppm belongs to the Sp stereoisomer.

The large difference in chemical shift for the U14 imino proton in the Rp isomer compared to the wild-type (~1.5 ppm) is in agreement with the different chemical nature of the hydrogen bond acceptor atom in the HB1 signature interaction, as detailed by the QM and QM/MM calculations (see the main text). In contrast, for the Sp isomer where the acceptor atom is oxygen, the chemical shift difference compared to the wild-type is rather small (less than 0.1 ppm). Assignments for the U13 and U18 imino proton signals to both stereoisomers in the chemically synthesized A17 PT were straightforward by observing the corresponding NOEs between the U14, U13 and U18 imino proton signals of each stereoisomer, respectively (Figure S10B).

*NMR experiments with the A16 PT NSR.*

Two sets of imino proton signals were also observed with the A16 PT variant for the U14 and U13 but not for U18 which yielded only a single signal (Figure 3C). Here, the PT influences the anion-π interaction (Figure 2) between the A16 phosphate group and the base of U14. In the Rp and Sp stereoisomers of A16 PT, sulfur and oxygen atoms, respectively, point toward the U14 base. The chemical shift difference between the two imino protons of U14 is much smaller than in the A17 PT variant (11.7 and 11.9 ppm) and closer to the wild-type (11.7 ppm). Importantly both U14 imino protons show correlations across hydrogen bond to a phosphate group at a chemical shift of ~ -3 ppm in the 2D-H,P long-range correlation experiment. This suggests that in both stereoisomers, the HB1 signature interaction with the A17 phosphate group is intact even with the A16 PT variant (Figure 3C). We assume that the signal with the chemical shift closer to the wild-type belongs to the Sp variant as the Rp variant should differ more from the WT. Therefore, we assign the signal at 11.9 ppm to the U14 imino resonance of the Rp stereoisomer. The assignments of the other signals can be derived again from a NOESY-spectrum (Figure S10C). Notably, the U13 imino proton signal assigned to the Rp isomer is shifted slightly upfield compared to the wild-type.

The assignment of the U13 and U14 imino resonances of the Rp stereoisomer is further supported by comparing the imino proton spectra of an A17C-mutant RNA with an A17C-mutant RNA transcribed in the presence of α-thio-ATP. The A17C-mutant RNA contains PT groups with the Rp configuration only at A16 and A24 while the A17C-mutant binds ribostamycin with the same binding mode as the wild-type albeit with a reduced affinity and adopts the same U-turn loop structure[3]. Comparison of the imino proton spectra of the A17C-mutant and the A16Rp_A17C_A24Rp RNA show that the A16Rp PT induces a slight downfield shift of the U14 imino proton (11.8 ppm). Furthermore, the U13 imino proton is shifted slightly upfield compared to the unmodified RNA (Figure S11A). These chemical shift changes have the same direction as those observed between the wild-type and the chemically synthesized A16Rp (Figure S11B).

*Details of NMR experiments with the C14 NSR containing the Rp stereoisomer of A17 PT.*

To gain more insights into the A17Rp/C14 PT structure, we transcribed the mutC14 RNA in the presence of α-thio-ATP and $^{13}C,^{15}N$-labelled CTP, yielding PT-modified RNA with Rp stereoisomer at nucleotides A16, A17 and A24. The protonation state of C14 in this RNA can be monitored in heteronuclear experiments using the $^{15}N$ and $^{13}C$-nuclei of the cytosine nucleotides[4] even when the H3 atom of the protonated C14 is not directly observable. In this case, we utilize $^{13}C$ chemical shift of the C4 carbon which can be observed in a 2D-H(N)C correlation experiment transferring magnetization from the amino groups to the C4.[5] The H(N)C-experiment recorded in our standard buffer at pH 6.2 for the C14 NSR with A16Rp, A17Rp, and A24Rp PTs shows a ~ 165.5 ppm chemical shift for the C4 carbon which is in the same range as the C4 carbon chemical shifts of all other C nucleotides in the RNA (Figure S9A). This confirms that unlike in the mutC14 system, the C14 is not protonated at pH 6.2 in the C14/A17Rp system. However, lowering the pH to 5.5 leads to a dramatic change in the C4 carbon chemical shift of C14 which is now observed at ~ 158 ppm (Figure S9A); a similar chemical shift as in the mutC14 systen at pH 6.2 (~ 158.5 ppm, Figure S9B) where C14 is stably protonated. The protonation is virtually complete at pH 5.5 as judged from the absence of line broadening, conforming that the $pK_a$ of C14 is still significantly higher than the $pK_a$ for free cytosine nucleotides (~ 4.5).

The $^{15}N$-HSQC spectra of the C14 NSR with A16Rp, A17Rp, and A24Rp PTs also show significant changes of the proton and nitrogen chemical shifts for the C14 amino group upon lowering the pH to 5.5 (Figure S9C), as expected from protonation of the neighboring N3 nitrogen. However, when the spectrum of the modified RNA at pH 5.5 is compared with the spectrum of the mutC14 RNA, it is striking that the N4 is shifted downfield by ~ 4 ppm in the modified RNA (~ 106 ppm) compared to the unmodified RNA (~ 102 ppm, Figure S9D) and that the chemical shifts of the two amino group protons show a much wider separation as well as a downfield chemical shift in the modified RNA. Similar observations were made for the chemical shifts in other systems when an oxygen was replaced by a sulfur as a hydrogen bond acceptor for an amino group. Thus, the data suggest that in presence of the Rp stereoisomer of A17 PT, the N4 amino group of the protonated C14 is involved in hydrogen bonding to the sulfur atom as well as the N3 imino group, leading to a bifurcated hydrogen bond. The presence of such a bifurcated hydrogen bond in the A17Rp/C14 RNA in contrast to the linear strong ionic hydrogen bond between the imino group and the phosphate in the mutC14 RNA might help to rationalize the lowered $pK_a$ for C14 protonation in the modified RNA.

*MD simulations and QM calculations of systems with Sp PT configuration.*

As mentioned in main text Table 1, we ran MD simulations on the systems with both Rp and Sp PT configurations. MD simulations of the systems with Sp configuration revealed no or only negligible differences among the key interactions of the NSR when compared to wild-type system. This included the key H-bonds, anion-π, and $K^+$ binding site. This result is not surprising as the force field describes the non-bridging oxygen atom using the same vdW parameter in both thiophosphate and phosphate, leading to virtually identical H-bond and anion-π geometries. Therefore, we do not present analyses of the MD simulation systems with Sp PT configuration in the main text.

The QM/MM and QM calculations, in principle, could be used to study the subtle differences between Sp PT and wild-type systems. However, interpretation of such differences is problematic when lacking the Boltzmann sampling of microstates obtained by the MD simulations, especially since both NMR and QM indicated the differences between wild-type and Sp PT NSR systems to be relatively minor. Rigorous examination of the Sp PT in NSR will require extensive experimental corroboration of the computational results and will be subject of a future study. In the present study, we show the Sp PT configuration data from the QM calculations only when such data is, in our opinion, conclusive and relevant for interpretation of the Rp PT data.

*Imperfect dimethyl-thiophosphate – water interaction energy scans.*

Comparison of QM and MM interaction energy scans for the dimethyl-thiophosphate (DMTP) – water system revealed that the QM optimized conformations are not always preferred by MM. We observed this for the DMTP – water interaction model, where the two water hydrogens in the complex were first QM refined, and the optimized complex then used for both QM and MM calculations. Such MM interaction energy curves, while physically relevant, do not fully represent the typical scenario encountered during simulations as QM minimum conformations might not be visited frequently in MD simulations.

Indeed, if we implement extra MM optimization on water hydrogens with both AMBER and thiolate parameters, the QM and MM curves will correspond to different conformations. Comparison of these two approaches is shown in Figure S4. After optimizing the complex in MM, the interaction energy in MM curves drops down significantly in terms of repulsion (SP2 – O(water) distance lower than 3.2 Å) albeit the optimum H-bond donor-acceptor distance does not have obvious change (Table S7). The different hydrogen orientations of water molecules in QM and MM optimized structures are further shown in the Supplementary Video. To briefly summarize, the QM optimized structures show clear repulsion between sulphur and the water hydrogens, while MM shows the water hydrogens pointing to the sulphur even if they are closer than 2 Å to each other. The vdW repulsion from sulphur to the water hydrogens is not captured by the optimization because there is no vdW sphere on hydrogens of the SPC/E water model we used.[6] This explains why the MM-optimized water hydrogens point to the sulphur even in the repulsive region, and why the interaction energy of the QM optimized structure is so high when computed by MM.

*Use of the HBfix protocol to optimize the description of the HB2 interaction.*

HBfix is an energy function added to the standard force field to tune the stability of selected hydrogen bonds.[7] It was originally designed to counteract the underestimation of base pair stability by introducing an extra potential on specific H-bonds. In our case, we applied the HBfix to improve the HB2 interaction in the NSR. The reversible transitions of the HB2 signature interaction between its native arrangement (Figure 2) and a non-native interaction with a nearby phosphate were indicated by a previous study.[8] The same behavior was observed in this study, for both the wild-type and PT simulations. As suggested by the previous study, we have tried to stabilize the HB2 by applying a 2 kcal/mol potential energy HBfix in a distance range of 2.3 Å and 3.3 Å between atoms U14(HO2') and A16(N7), thus supporting the native arrangement. Only thiolate sulphur parameters[9] were utilized in simulations where we applied HBfix. As shown in Table S8, the use of HBfix significantly improved the reproduction of the native HB2 in both the A17Rp/U14 and A16Rp/U14 systems with the populations of 97.4% (from 54.8%) and 87.0% (from 16.7%), respectively. In other words, HBfix solved the HB2 instability problem in NSR MD simulations.[8] Importantly, the HBfix produces this improvement without compromising other parts of the system in any way.

We also note the lower stability of the HB2 interaction in both the standard and HBfix simulations of the A16Rp/U14 system. This could suggest that the PT modification on A16 may be somewhat destabilizing for the HB2 interaction. However, due to the admittedly problematic description of the HB2 interaction by the force field, we refrain from making strong conclusion based on this observation.[8]

*Details of the interaction energy QM and MM scans.*

For the SP2 – H3-N3 H-bond model scan in DMTP - methyl-uracil (MU) pair, the increment was 0.05 Å and 0.1 Å from 2 Å to 3 Å and from 3 Å to 7 Å, respectively. For the OP2 – H3-N3 H-bond scan in dimethyl-phosphate (DMP) – MU pair, the increment was 0.05 Å and 0.1 Å from 1.4 Å to 3 Å and from 3 Å to 7 Å, respectively. The distance range of the SP2 – U14 base anion-π interaction model scan was from 2.5 Å to 4 Å and from 2.1 Å to 3.6 Å for OP2 – U14. The increment was 0.1 Å.

We also performed QM distance scans between DMP and a water molecule. The coordinates from A17Rp/U14 QM/MM optimized structure were used as the initial structure with the vector defined by the H3 – N3 bond of the extracted uracil corresponding to the hydrogen – oxygen bond of the water molecule with which we replaced the base (see Figure S3). The structure was used to obtain a series of structures with different OP2 – O or SP2 – O hydrogen bond distances. For each distance point, coordinates of the water hydrogens were optimized while all other atoms of the system were frozen. Afterwards, the QM and MM distance scan calculations were run analogously to the phosphate-base system described in the main text. The SP2 – O distance scan range was from 2.5 Å to 8 Å. The increment was 0.05 Å and 0.1 Å from 2.5 Å to 4 Å and 4 Å to 8 Å, respectively. The OP2 – O distance scan range was from 2.4 Å to 4 Å with increment of 0.05 Å.

Selected results obtained by PBEh-3c were further validated by the more rigorous DLPNO-CCSD(T)/TightPNO/CBS method[10,11]. The DLPNO-CCSD(T)/TightPNO/CBS method, used with aug-cc-pV(T-Q)Z and the auxiliary aug-cc-pVQZ/C basis sets along with TightSCF setting, was used as a reference QM method.[12–14] We applied this method for selected distance ranges of interest in both HB1 and anion-π QM interaction scans (Figure S2A, S5A). PBEh-3c gives slightly higher absolute interaction energy for HB1 than the more accurate QM methods, but the optimum HB1 distances and the overall curve tendencies among these methods are consistent. In anion-π scans, PBEh-3c and DLPNO-CCSD(T)/TightPNO/CBS curves essentially overlap. Overall, these results suggest that the performance of the PBEh-3c method is robust enough for use in our QM interaction energy scan study. Due to the utilized Lennard-Jones potentials,[15] the MM grossly overestimates the short-range repulsion of HB1 (1.2 to 2 Å for DMTP-MU and 1.0 to 1.5 Å for DMP-MU) and anion-π (2.0 to 2.7 Å for DMTP-MU and 1.5 to 2.2 Å for DMP-MU) compared to all the tested QM methods (Figure S2B and Figure S5B).

*Additional MD simulations with different sulphur parameters.*

The additional MD simulations of PT modified NSR were run either with global or NBfix thiolate parameters[9], which give the larger sulphur vdW radius as suggested by the QM and QM/MM calculations (see the main text). The populations of signature HB1 and HB2 interactions in the new simulations were consistent with the original simulations (Table S8). NBfix-implemented thiolate parameter in the A16Rp/U14 system (see the main text) provides the best description of the anion-π interaction as the weakened interaction with water, also in turn, somewhat stabilizes the anion-π interaction which is reflected in a narrower range of observed distances (Figure 12B) compared to the globally implemented AMBER parameter.

*Incomplete NBfix and its effects on solute-solvent interactions.*

In our initial parametrization attempts, termed as "incomplete NBfix", we only modified the sulphur non-bonded interactions with U14(H3) and water molecules. This resulted in imbalance of sulphur-ion and sulphur-water interactions. Specifically, the ion almost permanently attached to the A17(SP2) in A17Rp/U14 system, distorting the structure and rendering the remainder of the U-turn motif unstable (Figure S12A). In other words, the sulphur became too attractive for the ions compared to the water molecules, giving rise to a higher sulphur hydrophobicity (Figure S12B) and potentially disrupting the HB1 interaction. We subsequently added the U14(N3) and ion interactions into the NBfix implementation, as described in the main text. Thereby, we obtained a balanced description of solute-solvent interactions.

***Supplementary Figures and Tables:***

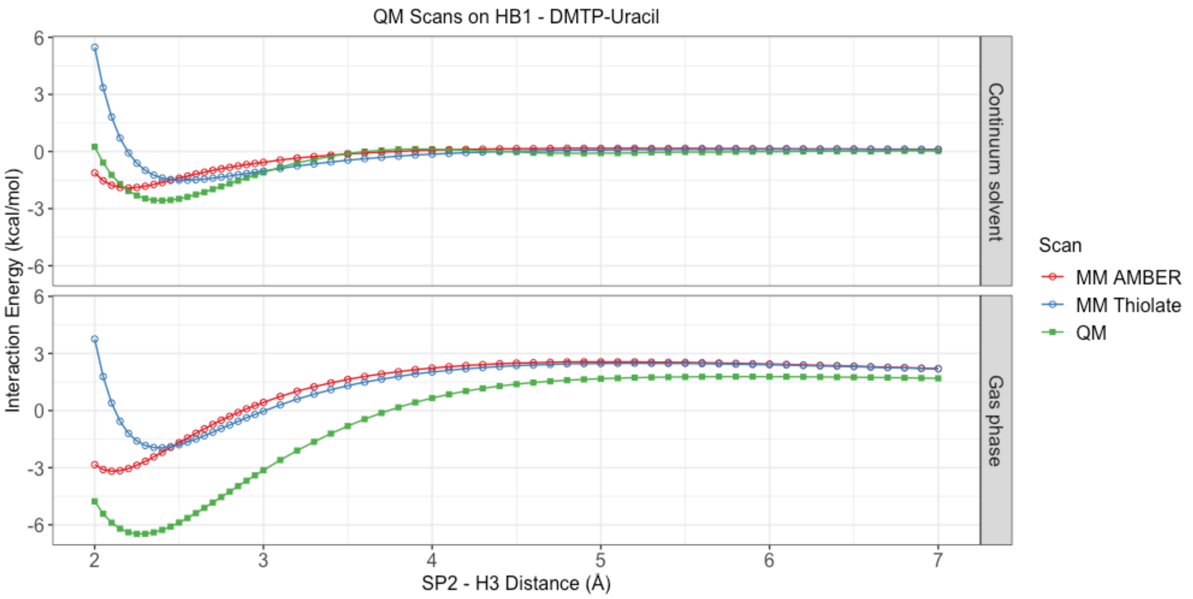


*Figure S1: QM and MM interaction energy scans for the HB1 interaction with the full distance range of the scan shown. Only the region of interest is shown in the main text figures.*

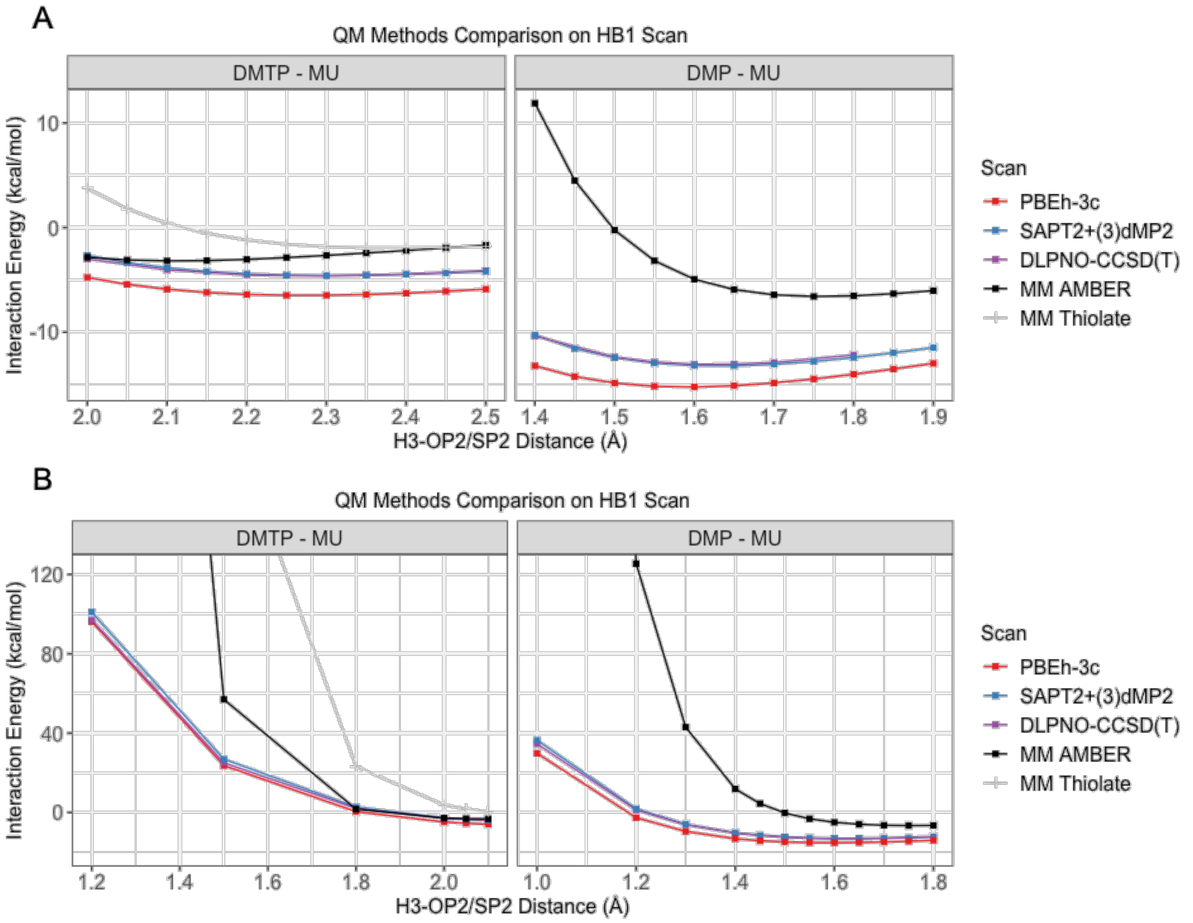


*Figure S2: A) Benchmark of the PBEh-3c method (used in the main text) against higher-level QM methods SAPT2+(3)δMP2, and DLPNO-CCSD(T), for QM distance scans of the HB1 interaction within a selected distance range. B) HB1 interaction scans in the short distance range.*

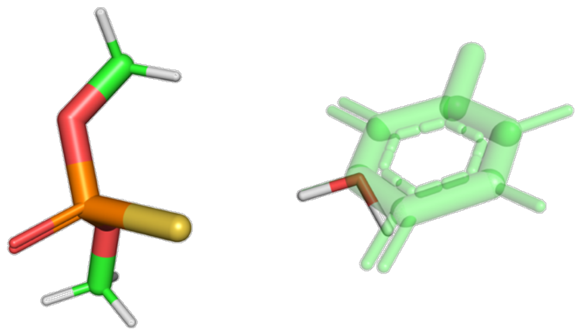

*Figure S3: Preparation of the small model system for the DMTP/DMP-water interaction for the H-bond scan study. The DMTP/DMP-water pair was derived from the QM/MM optimized DMTP/DMP-MU complex by fitting the water H-O bond to the H3-N3 bond. The uracil is in transparent green. It was not included in the calculation and is only visualized for the H3-N3 bond position.*

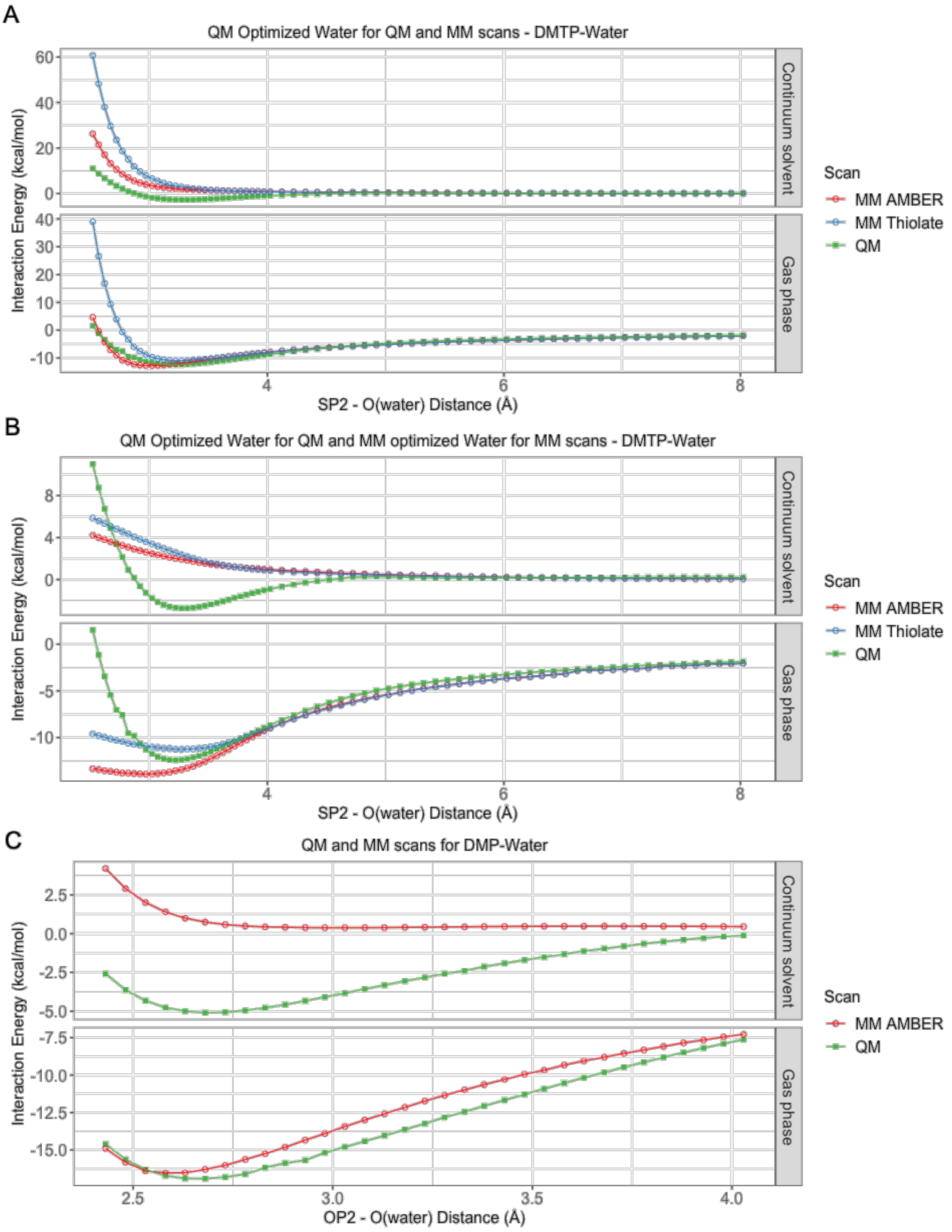


*Figure S4: QM and MM interaction energy scans against A) DMTP – water, where the orientations of water hydrogens were optimized by QM and then used in both QM and MM scans, B) the DMTP – water where the water hydrogen orientations in MM scans were further optimized by MM, and C) DMP – water, where the orientations of water hydrogens were optimized by QM and then used in both QM and MM scans.*

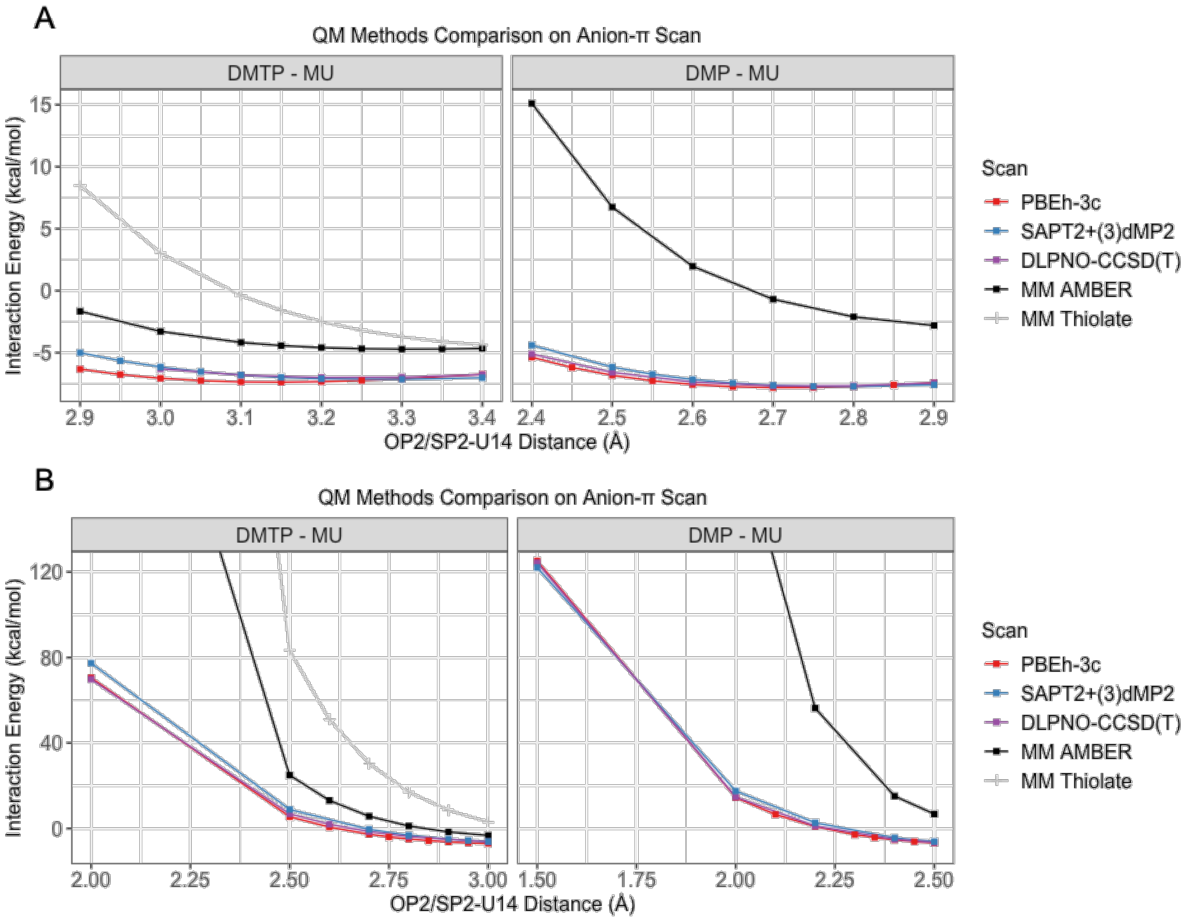


*Figure S5: A) Benchmark of PBEh-3c method (used in the main text) against higher-level QM methods, SAPT2+(3)δMP2 and DLPNO-CCSD(T)/CBS, for QM distance scans of the anion-π interaction, within the selected distance range. B) Anion-π interaction scans in the short distance range.*

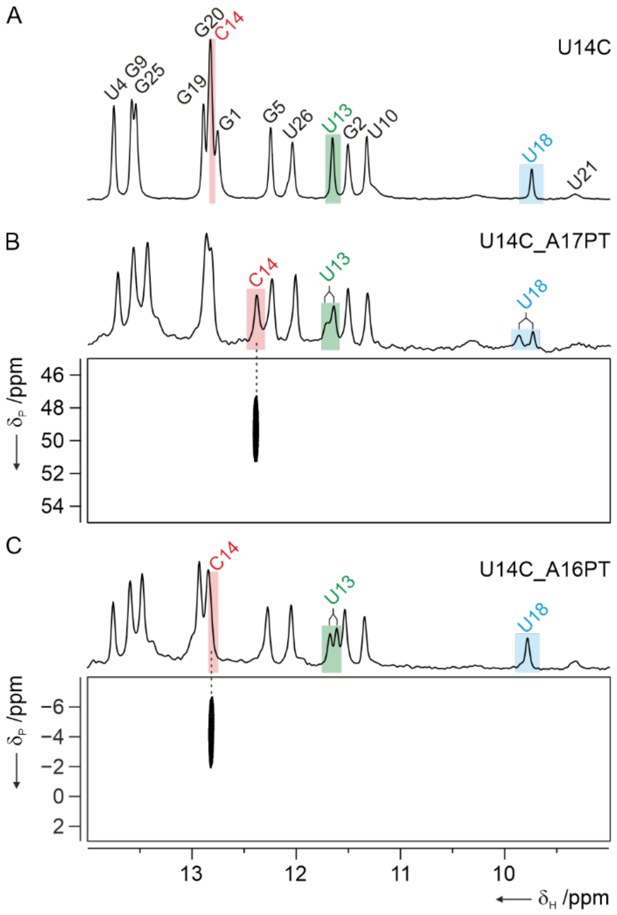


*Figure S6: Effect of phosphorothioate modifications on the structure of the U14C-NSR-RNA A)1D imino proton spectrum of the unmodified U14C-NSR bound to ribostamycin. Signal assignments are given and the assignments for the imino protons of the protonated C14, U13 and U18 are highlighted in different colors. B) Imino proton spectrum (top) and H,P-correlation experiment for the A17PT-modified U14C-RNA. C) Imino proton spectrum (top) and H,P-correlation experiment for the A16PT-modified U14C-RNA.*

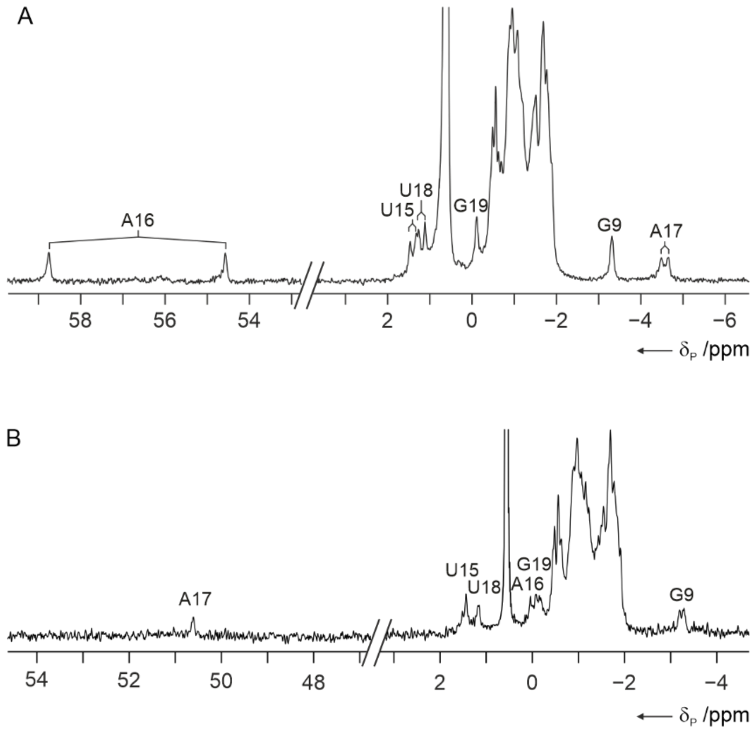


*Figure S7: $^{31}$P-NMR spectra of the A16PT and A17PT-modified U14C-NSR RNAs bound to ribostamycin at pH 6.2. A) $^{31}$P-NMR spectrum for the A16PT – U14C RNA. Signal assignments are indicated. B) $^{31}$P-NMR spectrum for the A17PT – U14C RNA.*

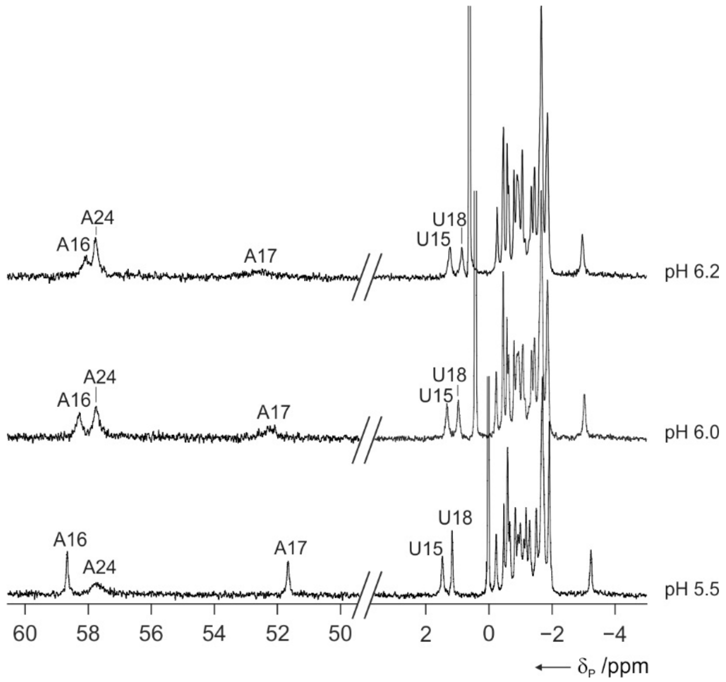


*Figure S8: Effects of pH on the 31P-1D-NMR spectra of the U14C_A16Rp_A17Rp _A24Rp-NSR RNA. Lowering of the buffer pH from pH 6.2 (top) to 5.5 (bottom) leads to the appearance of the signal for the A17 phosphorothioate group and to sharpening of the signals for U15, A16 and U18 in agreement with overall structure stabilization upon C14 protonation and a folding at low pH structurally similar to the one observed for the WT and the unmodified mutC14 RNA.*

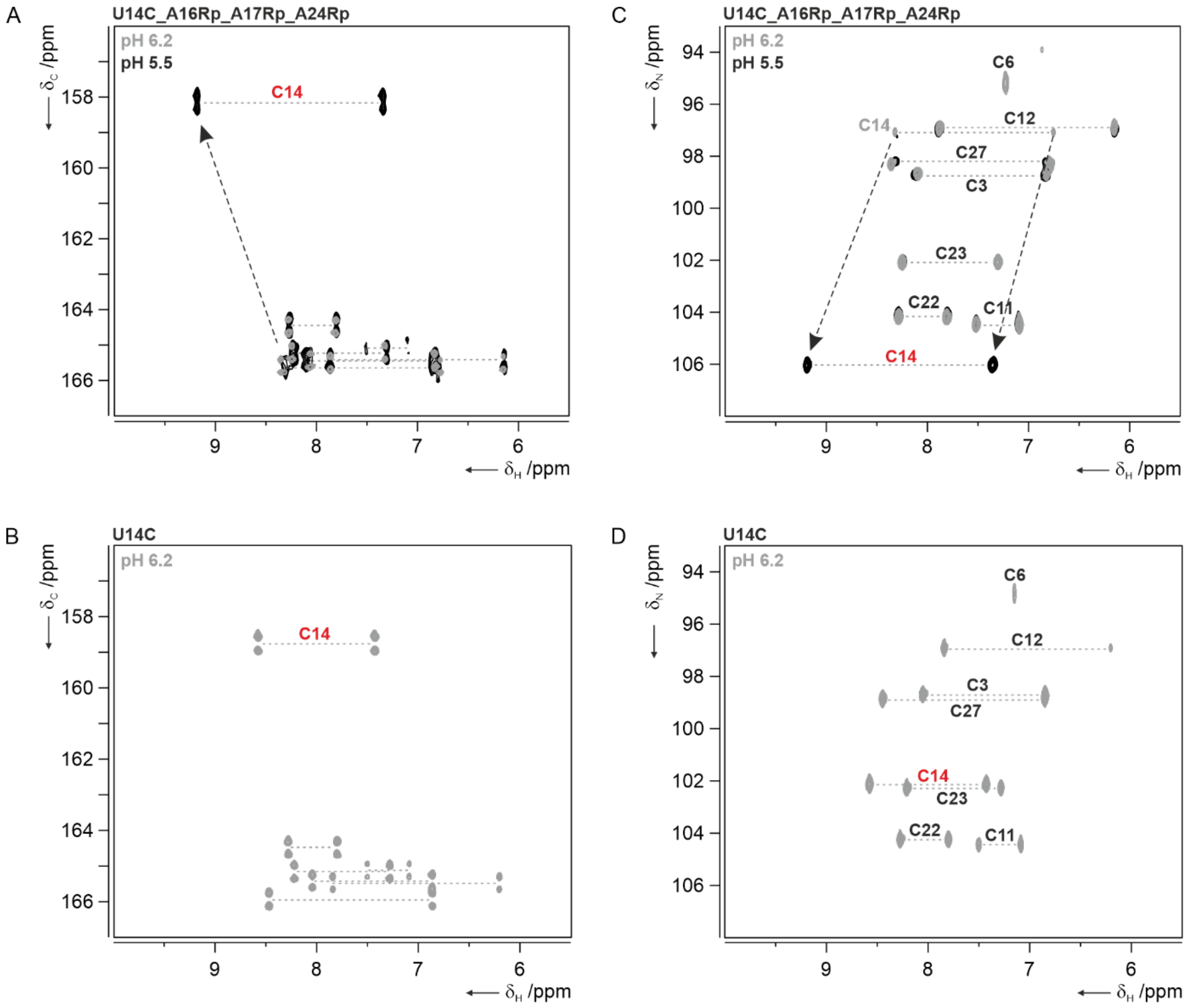


*Figure S9: Effects of pH on heteronuclear chemical shifts in the U14C_A16Rp_A17Rp _A24Rp-NSR RNA. A) Comparison of the 2D-H(N)C-spectrum recorded at pH 6.2 (grey) and 5.5 (black) for the modified RNA shows a dramatic change for the chemical shift of the C4 carbon of C14 upon lowering the pH. B) 2D-H(C)N-spectrum of the unmodified U14C-RNA at pH 6.2. The C4 carbon chemical shift of C14 in the stably protonated unmodified U14C-RNA at pH 6.2 is similar to the one measured for the modified RNA at pH 5.5 indicative of protonation. C) Comparison of 2D-$^{15}$N-HSQC-spectra for the C amino groups in the modified RNA at pH 6.2 (grey) and 5.5 (black). Assignments are given and the assignment for the C14 amino group is highlighted in red. Chemical shift changes are indicated by arrows. D) 2D-$^{15}$N-HSQC-spectrum for the C amino groups in the unmodified U14C-NSR RNA at pH 6.2. Assignments for individual amino groups are given and the assignment for the C14 amino group is highlighted in red.*

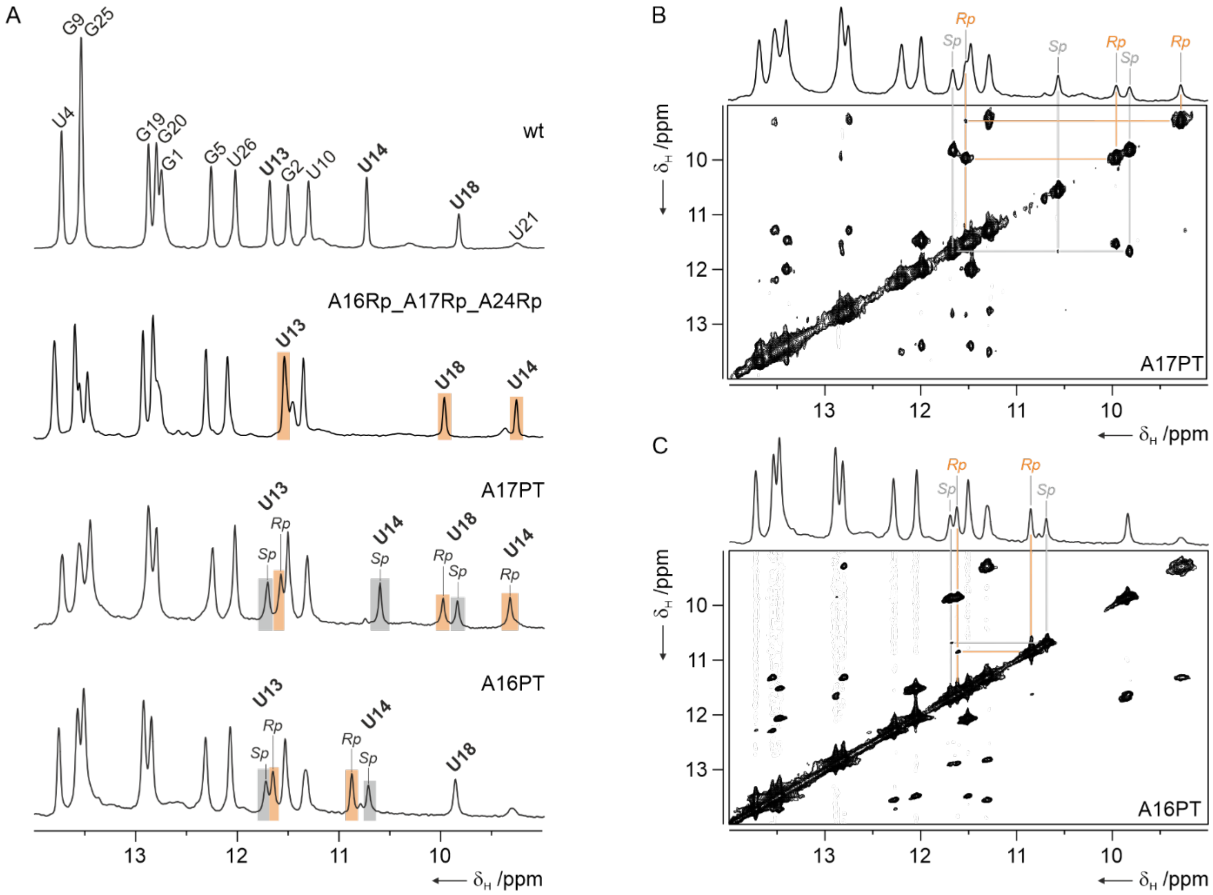


*Figure S10: Stereospecific signal assignments for the A17 PT and A16 PT modified RNAs. A) Comparison of the imino proton spectra of wild-type (top), A16 Rp, A17 Rp, A24 Rp PT substituted RNA prepared by in vitro transcription with T7-RNA-polymerase using α-thio-ATP instead of ATP (second from top), a chemically synthesized A17 PT modified RNA containing both the Rp and the Sp stereoisomer (second from bottom), and a chemically synthesized A16 PT modified RNA (bottom). Signals corresponding to the Rp and Sp stereoisomers are highlighted in orange and gray, respectively. All imino protons with the same chemical shift in both stereoisomers are unmarked. B) Imino proton region of a 2D-NOESY spectrum for the A17 PT NSR. C) Imino proton region of a 2D-NOESY spectrum for the A16 PT NSR. NOE-cross peaks connecting the U13, U14 and U18 imino protons in B and C are indicated and color-coded as in A.*

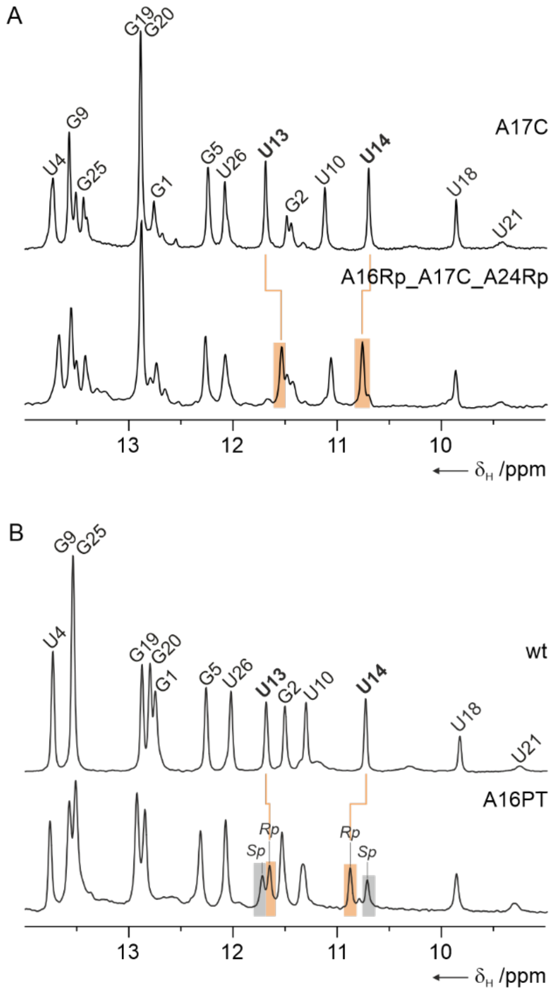


*Figure S11: Effects of A16Rp modification on chemical shifts in the A17C-mutant NSR RNA. A) Comparison of the imino proton spectra in the A17C (top) and A16Rp_A17C_A24Rp RNAs (bottom). The presence of the A16Rp PT leads to a downfield chemical shift change of the U14 imino proton and an upfield chemical shift change for the U13 imino proton. B) Chemical shift changes in the same directions are observed for the U13 and U14 imino proton signals of the wild-type NSR (top) and the putative Rp isomer in the A16 PT NSR (bottom).*

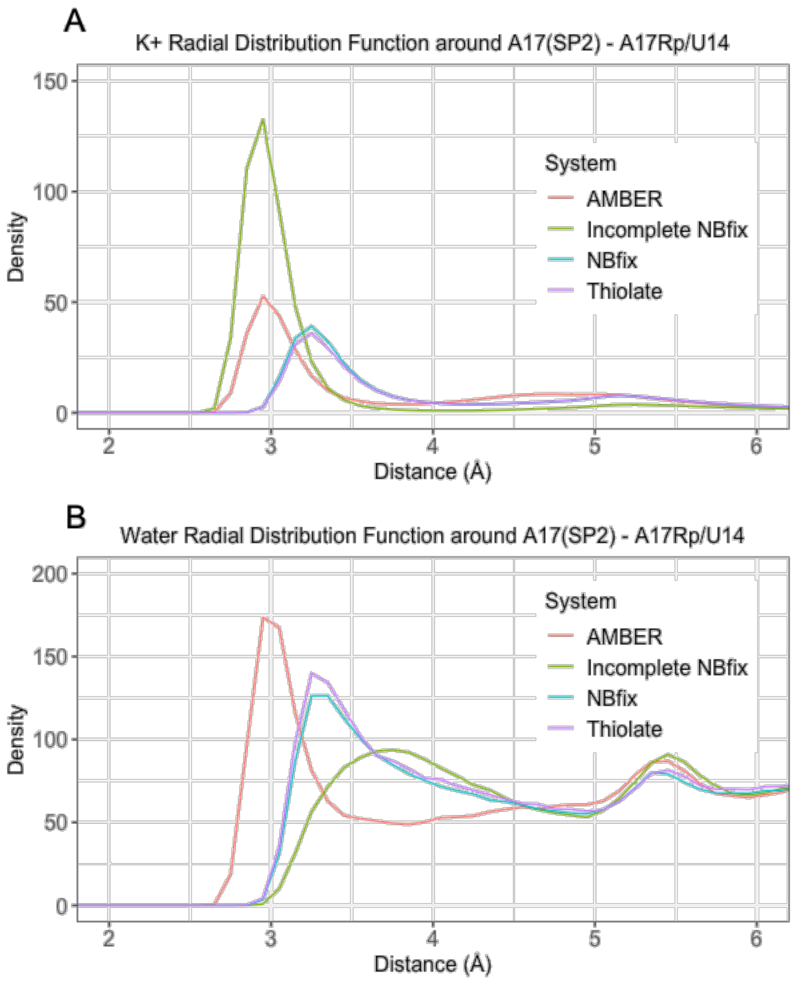


*Figure S12: A) Radial distribution functions for potassium ions in the vicinity of the A17(SP2) atom in the A17Rp/U14 simulations, with different parameter sets used. B) Radial distribution functions of water around A17(SP2) atom in the A17Rp/U14 simulations, with different parameter sets used.*

*Table S1: List of all MD simulations.*

| | simulated system | number of simulations × length (µs) |
|---|---|---|
| AMBER vdW Sulphur Parameters | A17Rp/U14<br>A17Sp/U14<br>A16Rp/U14<br>A16Sp/U14<br>A17Rp/C14<br>A17Sp/C14<br>A16Rp/C14<br>A16Sp/C14 | 4 × 1 |
| | wt<br>MutC14 | 1 × 10 |
| Other vdW Sulphur Parameters (either globally-implemented thiolate or NBfix) | A17Rp/U14 + thiolate<br>A16Rp/U14 + thiolate<br>A17Rp/U14 + NBfix<br>A16Rp/U14 + NBfix<br>A17Rp/U14 + incomplete NBfix<br>A17Rp/U14 + thiolate + HBfix (HB2)<br>A16Rp/U14 + thiolate + HBfix (HB2) | 4 × 1 |

*Table S2: The HB1 geometries in initial structures and after QM/MM and MM optimizations with AMBER parameters.*[a]

| HB1 (Å) | $D_0$(X-Y) | $D_{QM/MM}$(X-Y) | $D_{MM}$(X-Y) | $D_0$(H-Y) | $D_{QM/MM}$(H-Y) | $D_{MM}$(H-Y) | $D_0$(X-H) | $D_{QM/MM}$(X-H) | $D_{MM}$(X-H) |
|---|---|---|---|---|---|---|---|---|---|
| **A17Rp/U14** | **3.4529** | **3.3000** | **3.0652** | **2.4221** | **2.2920** | **2.0683** | **1.0472** | **1.0162** | **1.0059** |
| A17Sp/U14 | 2.7776 | 2.7789 | 2.7425 | 1.7435 | 1.7610 | 1.7524 | 1.0489 | 1.0313 | 1.0100 |
| **A17Rp/C14** | **3.1404** | **3.3028** | **3.0657** | **2.2130** | **2.3500** | **2.1380** | **1.0100** | **1.0252** | **1.0083** |
| A17Sp/C14 | 2.8203 | 2.7252 | 2.7957 | 1.7346 | 1.7015 | 1.8058 | 1.1006 | 1.0435 | 1.0136 |
| A16Rp/U14 | 3.3741 | 3.0096 | 2.8886 | 2.3776 | 1.9978 | 1.8806 | 1.0421 | 1.0235 | 1.0082 |
| A16Sp/U14 | 3.1024 | 2.9107 | 2.9084 | 2.1084 | 1.8992 | 1.9190 | 1.0407 | 1.0257 | 1.0114 |
| A16Rp/C14 | 2.9631 | 2.8529 | 2.8159 | 1.8983 | 1.8312 | 1.8087 | 1.0722 | 1.0314 | 1.0114 |
| A16Sp/C14 | 2.9353 | 2.7858 | 2.8415 | 1.8927 | 1.7476 | 1.8267 | 1.0494 | 1.0441 | 1.0165 |
| wt | 2.7695 | 2.8829 | 2.7780 | 1.7065 | 1.8646 | 1.7850 | 1.0885 | 1.0262 | 1.0120 |
| mutC14 | 2.7720 | 2.6734 | 2.7388 | 1.7400 | 1.6470 | 1.7540 | 1.0678 | 1.0457 | 1.0125 |

[a] *The table shows the HB1 distance (D) comparisons among the starting (0), QM/MM and MM optimized structures. "X" is the donor, "Y" is the acceptor, and "H" is the hydrogen within the H-bond. Difference of the QM/MM and MM optimized structures is the key parameter to follow irrespective of the value of the starting structure.*

*Table S3: Comparison of the HB1 and HB1a distances and angles in the initial structures and in the QM/MM and MM optimized systems. The data in this table refer to the same structures as in Table S2.*

| H-Bond Length (Å) | A17(OP/SP) - H3 | | | A17(OP/SP) - H41 | | |
|---|---|---|---|---|---|---|
| | Initial | QM/MM | MM | Initial | QM/MM | MM |
| A17Rp/C14 | 2.2130 | 2.3503 | 2.1380 | 2.2362 | 2.5395 | 2.2145 |
| A17Sp/C14 | 1.8286 | 1.8139 | 1.7794 | 1.9575 | 1.9158 | 2.0042 |
| A16Rp/C14 | 1.8721 | 1.7995 | 1.7661 | 1.7722 | 2.3309 | 1.8745 |
| A16Sp/C14 | 1.8795 | 1.6084 | 1.8404 | 1.9340 | 2.9011 | 1.7993 |
| mutC14 | 1.8340 | 1.6469 | 1.7540 | 2.2273 | 2.6073 | 2.5496 |

| | | | | | | |
|---|---|---|---|---|---|---|
| mutC14 from A17Sp/C14[a] | 1.8286 | 1.7596 | 1.7812 | 1.9575 | 2.1242 | 1.8752 |

| H-Bond Angle (º) | A17(OP/SP) – C14(H3) – C14(N3) | | | A17(OP/SP) – C14(H41) – C14(N4) | | |
|---|---|---|---|---|---|---|
| | Initial | QM/MM | MM | Initial | QM/MM | MM |
| A17Rp/C14 | 151.9 | 154.1 | 152.1 | 155.8 | 144.0 | 149.1 |
| A17Sp/C14 | 145.6 | 147.7 | 154.3 | 144.8 | 141.0 | 145.3 |
| A16Rp/C14 | 146.5 | 159.5 | 152.1 | 152.2 | 135.4 | 147.7 |
| A16Sp/C14 | 144.4 | 168.0 | 148.7 | 142.4 | 116.3 | 151.2 |
| mutC14 | 163.4 | 165.9 | 163.1 | 145.7 | 123.2 | 129.9 |
| mutC14 from A17Sp/C14[a] | 145.6182 | 154.5243 | 151.2256 | 144.8318 | 137.7753 | 148.4457 |

[a]*Starting structure for the* "mutC14 from A17Sp/C14" *system was derived from optimized A17Sp/C14 structure by replacing the sulphur with oxygen.*

*Table S4: SAPT values calculated at optimum acceptor-hydrogen distances for HB1 (2.30 Å and 1.63 Å for DMTP-MU and DMP-MU, respectively).*

| *Type* | *DMTP-MU* | | *DMP-MU* | |
|---|---|---|---|---|
| | *E / (kcal/mol)* | *%[a]* | *E / (kcal/mol)* | *%* |
| *Total* | -4.58 | | -13.22 | |
| *Disp* | -5.40 | 34 | -8.15 | 21 |
| *Elec* | -4.14 | 26 | -16.85 | 42 |
| *Exch* | 11.21 | | 26.24 | |
| *Ind* | -6.26 | 40 | -14.46 | 37 |

[a] *Only the negative energy terms, namely dispersion, electrostatics and induction, show the percentage value, which is calculated over the sum of these energy terms.*

*Table S5: Summary of the QM and MM HB1 scans on DMTP/DMP – water.*

| System | Calculation method | Optimum distance (Å)[a] | Minimum energy (kcal/mol) |
|---|---|---|---|
| DMTP – water | | | |
| | MM AMBER / vacuum | 3.01 | -12.79 |
| | MM thiolate / vacuum | 3.25 | -10.99 |
| | MM AMBER / GBSA | -[b] | -[b] |
| | MM thiolate / GBSA | -[b] | -[b] |
| | QM / vacuum | 3.22 | -12.40 |
| | QM / COSMO | 3.29 | -2.87 |
| DMP – water: | | | |
| | MM AMBER / vacuum | 2.61 | -16.54 |
| | MM AMBER / GBSA | -[b] | -[b] |
| | QM / vacuum | 2.66 | -16.93 |
| | QM / COSMO | 2.69 | -5.21 |

[a] *The distance between sulphur (or oxygen in DMP) and water oxygen which is associated with the lowest interaction energy.* [b]*The optimum distance was not found within the scanned distance range. More details of the scans are in SI Appendix.*

*Table S6: SAPT values calculated at optimum distances for anion-π interaction (3.27 Å and 2.79 Å for DMTP-MU and DMP-MU, respectively)*

| *Type* | *DMTP-MU* | | *DMP-MU* | |
|---|---|---|---|---|
| | *E / (kcal/mol)* | *%* | *E / (kcal/mol)* | *%* |
| *Total* | -7.16 | | -7.73 | |
| *Disp* | -7.08 | 42 | -7.12 | 41 |
| *Elec* | -6.73 | 40 | -4.88 | 28 |
| *Exch* | 9.79 | | 9.47 | |
| *Ind* | -3.14 | 19 | -5.21 | 30 |

*Table S7: Comparison of the DMTP-water MM scans with water molecules optimized by QM and MM methods.*

| Water optimization methods | Calculation method | Optimum distance (Å) | Minimum energy (kcal/mol) |
|---|---|---|---|
| QM: | | | |
| | MM AMBER / vacuum | 3.01 | -12.79 |
| | MM thiolate / vacuum | 3.25 | -10.99 |
| | QM / vacuum | 3.22 | -12.40 |
| MM: | | | |
| | MM AMBER / vacuum | 2.95 | -13.91 |
| | MM thiolate / vacuum | 3.23 | -11.27 |

*Table S8: Populations of HB1 and HB2 signature H-bonds in MD simulations.*

| Systems | HB1 Population (%) | HB2 Population (%) |
|---|---|---|
| A17Rp/U14 | 93.9 | 54.8 |
| A16Rp/U14 | 94.5 | 16.7 |
| A17Rp/U14 incomplete NBfix | 96.7 | 45.6 |
| A17Rp/U14 NBfix | 88.1 | 42.0 |
| A16Rp/U14 NBfix | 98.0 | 30.8 |
| A17Rp/U14 thiolate | 95.8 | 61.2 |
| A16Rp/U14 thiolate | 94.0 | 22.0 |
| A17Rp/U14 HBfix | 98.5 | 97.4 |
| A16Rp/U14 HBfix | 85.3 | 87.0 |
